# Lactate-carried Mitochondrial Energy Overflow

**DOI:** 10.1101/2024.07.19.604361

**Authors:** Daniela Rauseo, Yasna Contreras-Baeza, Hugo Faurand, Nataly Cárcamo, Raibel Suárez, Alexandra von Faber-Castell, Franco Silva, Valentina Mora-González, Matthias T. Wyss, Felipe Baeza-Lehnert, Iván Ruminot, Carlos Alvarez-Navarro, Alejandro San Martín, Bruno Weber, Pamela Y. Sandoval, L. Felipe Barros

## Abstract

We addressed the question of mitochondrial lactate metabolism using genetically-encoded sensors. The organelle was found to contain a dynamic lactate pool that leads to dose- and time-dependent protein lactylation. In neurons, mitochondrial lactate reported blood lactate levels with high fidelity. The exchange of lactate across the inner mitochondrial membrane was found to be mediated by a high affinity H^+^-coupled transport system involving the mitochondrial pyruvate carrier MPC. Assessment of electron transport chain activity and determination of lactate flux showed that mitochondria are tonic lactate producers, a phenomenon driven by energization and stimulated by hypoxia. We conclude that an overflow mechanism caps the redox level of mitochondria, while saving energy in the form of lactate.

**One Sentence Summary:** Mitochondrial lactate production

## Main Text

Mitochondria are energized by pyruvate, which is generated in the cytosol from glucose or lactate. Once in the mitochondrial matrix, pyruvate is oxidized to CO_2_ via the Krebs cycle, generating NADH and FADH_2_, which are used by the electron transport chain (ETC) to pump protons towards the mitochondrial intermembrane space. This flux of protons builds the electromotive force that drives the production of ATP by oxidative phosphorylation (OxPhos). Additional OxPhos substrates are fatty acids, amino acids and ketone bodies, all of which energize mitochondria via the ETC.

Lactate, which is made from pyruvate by the redox enzyme lactate dehydrogenase (LDH), serves to exchange carbons and energy between cells and tissues passing through monocarboxylate transporters (MCTs; *1-4*). Lactate is also an intercellular signal, acting through various mechanisms including redox ratio, G protein-coupled receptors, and post-translational modifications (*5-8*). In addition to these roles, lactate has been proposed to be directly oxidized by mitochondria, a model known as the Intracellular Lactate Shuttle (ILS; *9, 10*). Despite receiving considerable experimental attention, ILS has remained controversial (*4, 10*), partly because of contrasting results and technical issues regarding purity of subcellular fractions, specificity of immunohistochemistry, and poor retention of radiolabeled metabolites. Here we have approached the issue of mitochondrial lactate metabolism using genetically encoded fluorescent indicators.

## RESULTS

Most experiments were performed in HEK293 cells, a relatively oxidative non-transformed epithelial cell line that is easy to handle. Other cell lines, brain cells in culture, and neurons *in vivo* were studied to ascertain the general applicability of the findings.

### A dynamic lactate pool within mitochondria

According to the ILS hypothesis there is lactate within mitochondria. In order to look for it, the single-fluorophore lactate indicator CanlonicSF (*11*) was stably expressed in HEK293 cells (HEK040 cell line), where it displayed a typical mitochondrial pattern, and showed co-localization with the voltage-sensitive mitochondrial marker TMRM (fig. S1). In the presence of physiological substrates, a steady-state pool of 1.2 mM lactate was found within the mitochondrial matrix (n=5; 53 cells; Fig. 1A-B; fig. S2-S3). Determined in other cell types, mitochondrial lactate levels ranged between 0.6 and 1.5 mM (Fig.1B; fig. S3), suggesting that the pool is conserved. In response to extracellular glucose or lactate, mitochondrial lactate increased in the order of seconds (Fig.1Cs; fig. S4). Cytosolic and mitochondrial lactate moved in register, indicating that the permeability of the mitochondrial membrane is in the same order of magnitude as that of the plasma membrane. CanlonicSF is affected by Ca^2+^ in the supraphysiological range (*11*), but control experiments showed that cytosolic and mitochondrial Ca^2+^ levels do not rise in response to lactate (fig. S5). Lactate regulates histones and other proteins by a post-translational modification termed lactylation (*7, 12*). Figure 1D and fig. S6 show that lactate exposure lead to widespread lactylation of HEK293 proteins in a dose- and time-dependent fashion. Several mitochondrial proteins became modified, including malate dehydrogenase (MDHM), lactate dehydrogenase B (LDHB) and superoxide dismutase (SODC; Figs. 1E-F; figs. S7-S8). These modifications are of potential functional interest as MDHM is part of the malate aspartate shuttle (MAS), and LDH and SODC are involved in hypoxia, see below.

**Fig. 1.**
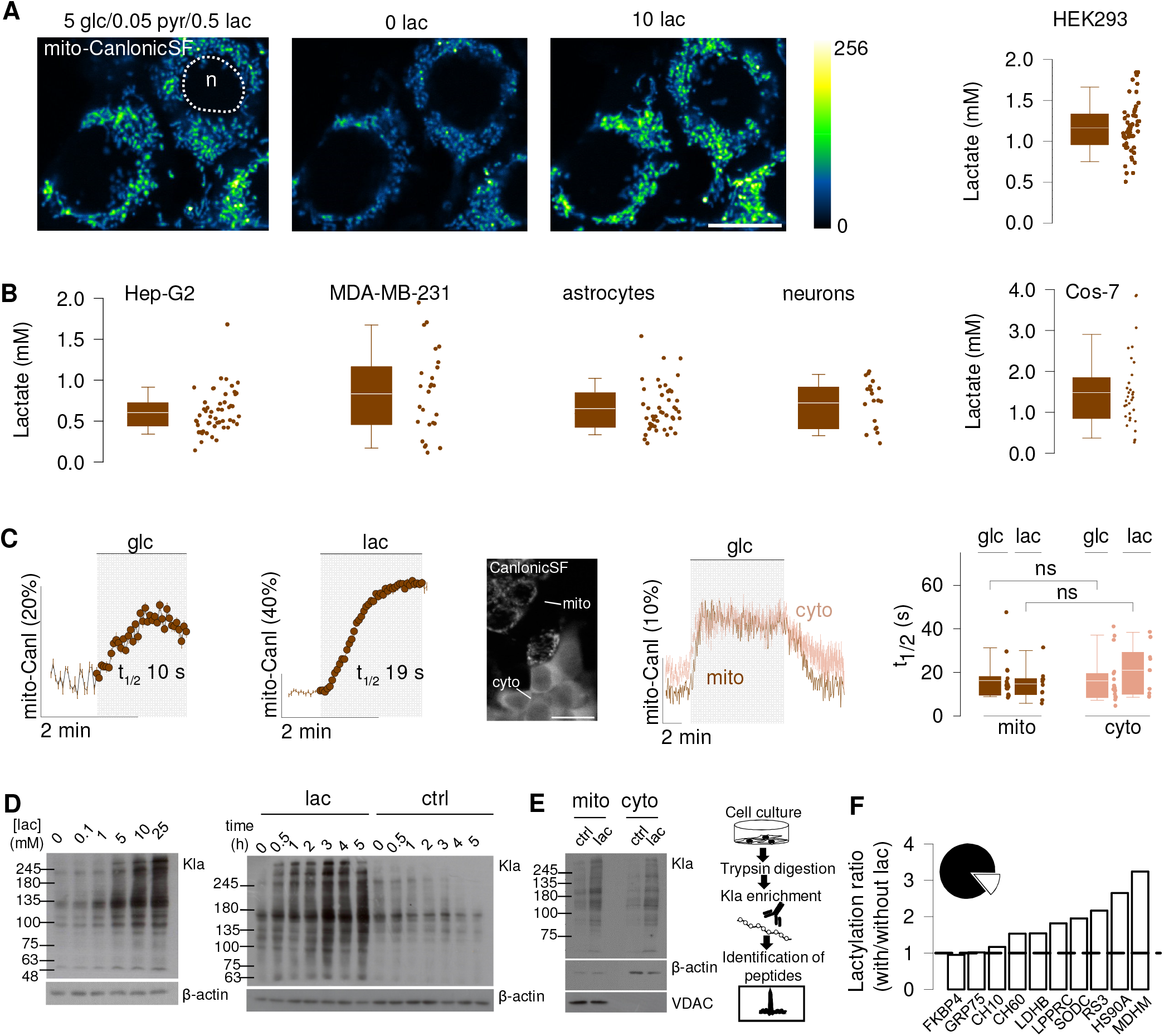
A dynamic mitochondrial lactate pool leads to protein lactylation. (**A**) HEK293 cells stably expressing mitoCanlonicSF (HEK040) were sequentially imaged at 5 mM glucose/0.05 mM pyruvate/0.5 mM lactate (standard buffer), 6 mM oxamate (nominal zero lactate) and high lactate (10 mM). Bar represents 20 µm. Summary of lactate levels in standard buffer (five experiments, 53 cells). (**B**)The same protocol was applied to: HepG2 (three experiments, 51 cells), MDA-MB-231 (three experiments, 26 cells), astrocytes (eleven experiments, 45 cells), neurons (eleven experiments, 19 cells) and Cos7 cells (six, 32 cells). For brain cells, the standard buffer contained 2 mM glucose/0.05 mM pyruvate/0.5 mM lactate.(**C**) Effect of glucose (5 mM) or lactate (10 mM) on HEK293 cells expressing mitoCanlonicSF or cytosolic CanlonicSF, measured simultaneously. Box plot shows data from three experiments, 18 cells (Mann-Whitney, paired t-test). Bar represents 20 µm. NS, not significant.(**D**) Lactate dose- and time-response of whole-cell HEK293 protein lactylation (Kla antisera). (**E**) Effect of lactate (25 mM/5 hours) on protein lactylation (Kla antisera) of HEK293 subcellular fractions. Preparation purity was assessed by detection of VDAC (mitochondria) or β-actin (cytosol). (**F**) Effect of lactate (25 mM/5 hours) on the lactylation level of HEK293 mitochondrial proteins, identified by nHPLC-MS/MS. Inset shows that 13% of precipitated proteins were mitochondrial. NS, not significant.

To look for the mitochondrial lactate pool *in vivo*, mito-CanlonicSF was targeted to mouse somatosensory cortex neurons and imaged by two-photon microscopy (Fig. 2A-B). An intravenous bolus injection of lactate led to a rise in blood lactate that peaked at 6 mM, a level reported in exercising human subjects (Fig. 2C; *13*). As shown in Figs. 2D-E, this perturbation caused a rise in mitochondrial lactate that closely followed cytosolic lactate, as detected with Laconic (*14, 15*). Thus, neuronal mitochondria are capable of sensing physiological fluctuations in blood lactate.

**Fig. 2.**
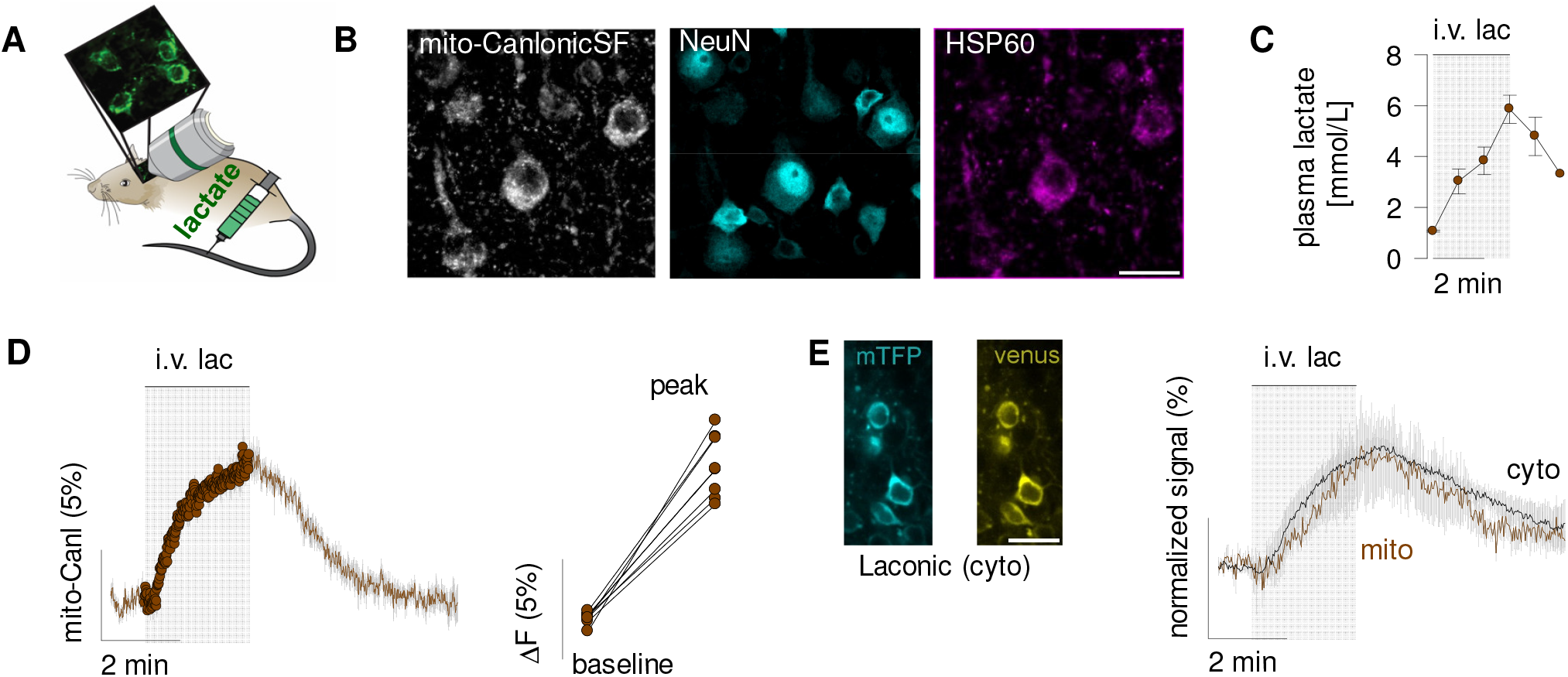
Mitochondrial lactate in neurons covaries with plasma lactate. (**A**) Mice expressing mito-CanlonicSF in neurons were i.v. injected with lactate (1.5 mmol/kg bodyweight). (**B**) Colocalization of mito-CanlonicSF with the neuronal marker NeuN and mitochondrial marker HSP60. Bar represents 20 µm. (**C**) Blood plasma lactate measurements upon lactate injection (data from three experiments in two animals). (**D**) Mito-CanlonicSF response upon lactate injection (data from eight experiments in four animals). (**E**) Paired measurement of lactate in mitochondria and cytosol, with mito-CanlonicSF and Laconic, respectively (normalized to peak, data from three experiments in three animals).

### Mitochondria transport lactate

Direct access to mitochondria was achieved by permeabilizing the plasma membrane (Fig. 3A), a treatment that preserves mitochondrial respiration (*8, 16*). The process was monitored by the release of cytosolic dyes and proteins (fig. S9A-B). Integrity of the outer mitochondrial membrane was corroborated by retention of a fluorescent protein targeted to the intermembrane space (fig. S9C), and by a functional ETC (see below), which requires retention of cytochrome C in the intermembrane space. In intact cells, exposure to pyruvate led to mitochondrial matrix acidification, explained by the entry of protons to the cytosol mediated by MCTs (figs. S9D-F; *17*). In contrast, in permeabilized cells, exposure to pyruvate caused mitochondrial matrix alkalinization caused by ETC activity (see below). The response to pyruvate was thus exploited as a routine demonstration of the quality of the preparation.

**Fig. 3.**
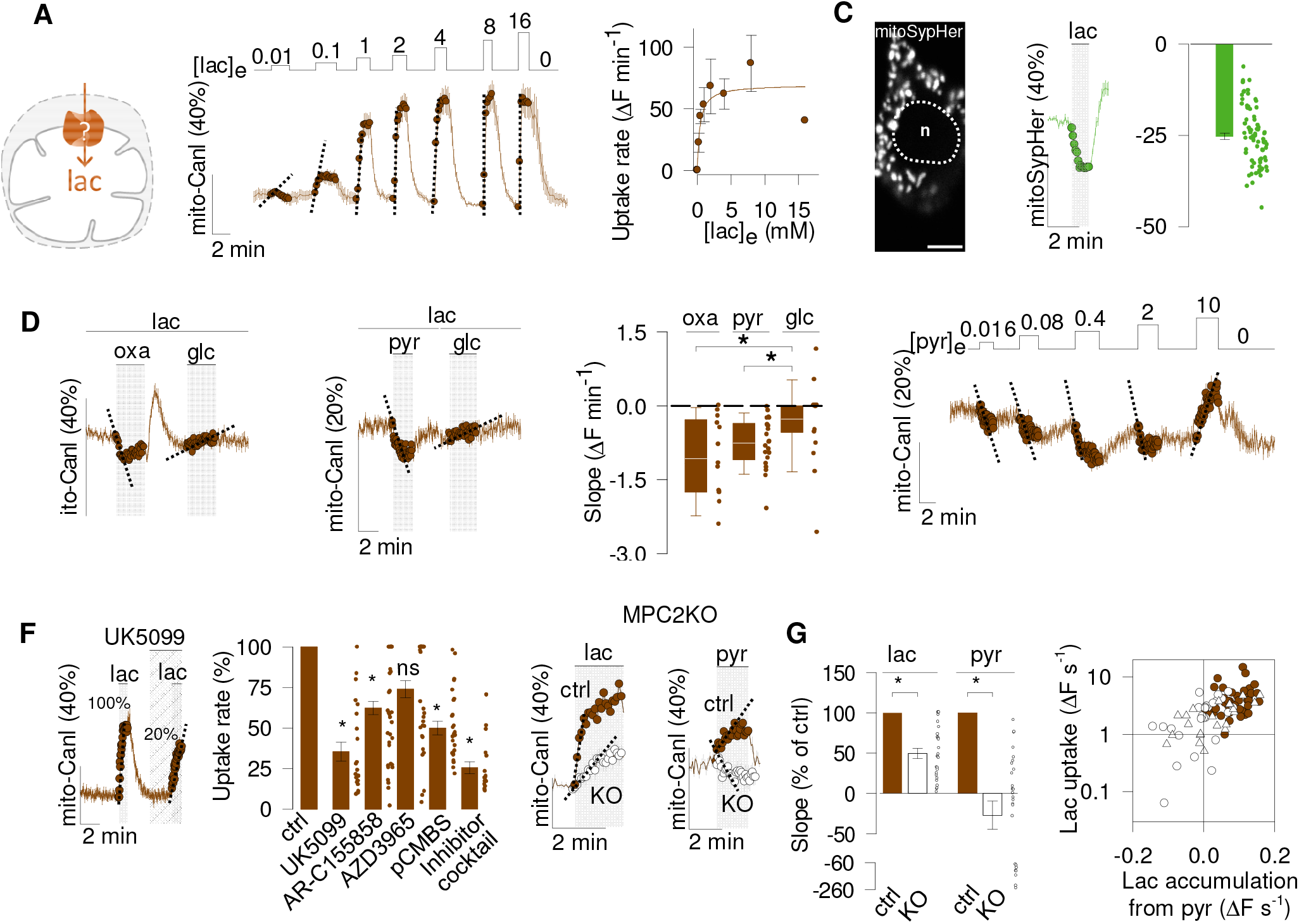
Characterization of mitochondrial lactate transport. Direct access of lactate to mitochondria was obtained by permeabilizing HEK293 cells expressing mito-CanlonicSF or mitoSypHer with digitonin (**A**). Except for the lactate dose response curve, mitochondria were kept energized with 0.2 mM glutamate plus 0.1mM malate. (**B**) Dose response of lactate uptake (mM), in cells expressing mito-CanlonicSF. A rectangular hyperbola was fitted to the initial uptake rates giving a K_M_= 0.8 ± 0.3 (four experiments, 75 cells). (**C**) Cells expressing the pH sensor mitoSypHer. The nucleus is indicated (n). Bar represents 20 µm. Effect of 10 mM lactate on matrix pH. Bar graph summarizes data from fifteen experiments (76 cells). (**D**) Mitochondria perfused with 0.5 mM lactate were exposed to oxamate (6 mM), pyruvate (0.5 mM) and glucose (5 mM). Initial slopes of lactate depletion are shown (two experiments,13 cells, paired t-test). (**E**) Mitochondria perfused with 0.5 mM lactate were exposed to increasing concentrations of pyruvate (mM; two experiments, 19 cells). (**F**) Lactate uptake was measured in the absence and presence of UK5099 (0.5 µM). Bar graph summarizes data from similar experiments with a panel of inhibitors. Ctrl (DMSO 0.01%-0.05%; nine experiments, 57 cells); UK5099 (0.5 μM; four experiments, 23 cells); AR-C155858 (1 μM; five experiments, 38 cells); AZD3965 (10 μM; five experiments, 36 cells); pCMBS (50 μM; four experiments, 25 cells); Inhibitor cocktail contains UK5099, AR-C155858, pCMBS and 5 μM syrosingopine (three experiments, 18 cells). Paired t-test. (**G**) Cells genetically depleted of MPC2 (sgMPC2KO7) were sequentially exposed to lactate (10 mM) or pyruvate (10 mM). Bar graph shown the summary of three KO experiments (27 cells) and three control experiments (40 cells). Student’
ss t-test. The right panel shows the cell-to-cell correlation between the initial rate of lactate production from pyruvate, and the rate of lactate uptake (•, ctrl; ○, sgMPC2KO7). Data obtained with a second RNA guide are also shown (Δ, sgMPC2KO9, three experiments, 22 cells).

The uptake of lactate by mitochondria was saturable, displayed a Michaelis-Menten constant K_M_ of 0.8 mM (Fig. 3B), and caused matrix acidification (Fig. 3C), pointing to a H^+^-coupled mechanism. Added to lactate-loaded mitochondria, the non-metabolized pyruvate analog oxamate was able to elicit the phenomenon of trans-acceleration, characterized by depletion followed by an overshoot upon substrate removal (Fig. 3D). At physiological concentrations pyruvate also trans-accelerated lactate, showing that the mitochondrial lactate transport system also carries pyruvate. Surprisingly, at higher pyruvate concentrations the depletion of lactate reverted into accumulation (Fig. 3E). Production of lactate from pyruvate pointed to LDH activity, a key observation that was pursued below.

The possibility that MCTs are involved in mitochondrial lactate transport was approached by pharmacological means. Experimental conditions were set so that two consecutive lactate pulses resulted in similar uptake rates (fig. S10). The potent MCT1 and MCT2 blocker AR-C155858 (*18*), which inhibits surface monocarboxylate uptake in HEK293 cells by > 95% (*15, 19, 20*), was very effective at preventing lactate entering mitochondria in intact cells (fig. S11). However, assayed in permeabilized cells, the inhibition of lactate uptake was much weaker (fig. S12). AZD3965, another MCT1/MCT2 blocker (*21*), was also relatively ineffective. pCMBS, a blocker of MCT1 and MCT4, but not of MCT2 (*22*), inhibited mitochondrial lactate uptake by about 50% (Fig. 3F; fig. S12). Further evidence against necessary MCT1 involvement was provided by the observation of robust mitochondrial uptake in MDA-MB-231 (fig. S13), a cell line that is genetically devoid of MCT1 (*19, 23*). In MDA-MB-231 mitochondria, uptake inhibition by syrosingopine, which targets MCT1 and MCT4, was also partial (fig. S13). To our surprise, the most effective inhibitor of mitochondrial lactate uptake was UK-5099 (Fig. 3F, fig. S12), a blocker of the mitochondrial pyruvate carrier (MPC; *22, 24, 25*). When lactate uptake was probed in the presence of a cocktail containing AR-C155858, pCMBS, syrosingopine and UK-5099, the inhibition was not stronger than that obtained with UK-5099 alone. In short, the functional and pharmacological profiles of mitochondrial lactate transport do not match that of typical plasma membrane MCTs. In view of the effectiveness of UK-5099, we probed cells devoid of MPC2 using CRISPR (*26*), a manipulation that abolished the entry of pyruvate (fig. S14). MPC2 deletion reduced the uptake of lactate by about 50% (Fig. 3G; fig. S15). These pharmacological and genetic interventions converge to indicate that the MPC plays a role in the transport of lactate. Of note, MPC deletion curtailed the production of lactate from pyruvate (Fig. 3G; fig. S15), providing independent support to the conclusion that there is LDH activity in the mitochondrial matrix.

### Mitochondria can consume lactate

Having demonstrated inner-membrane lactate transport and matrix LDH activity, the two pillars of ILS, we sought proof of effective lactate metabolism using autofluorescence, which reflects the redox status of the mitochondrial matrix (*27-29*). Exposure to pyruvate or engaging the MAS with glutamate/malate caused a small decrease in mitochondrial autofluorescence (Fig. 4A), but lactate was ineffective. A stronger change was obtained by applying pyruvate after pre-energization with glutamate/malate, but lactate was still without effect (Fig. 4A). In principle, the generation of reduced cofactors could be matched by their oxidation, making autofluorescence a poor proxy of metabolism. Thus, we developed a more sensitive method to detect energization, by imaging matrix pH. The rationale of this approach is that if the F_1_F_o_-ATPase is inactive, ETC activity can be made visible as matrix alkalinization, equivalent to State 3 respiration (Fig. 4B; *29*). Indeed, addition of pyruvate elicited a rapid and sustained rise in matrix pH, which was precluded by pharmacological ETC inhibition (Fig. 4C-D; fig. S16). Mimicking the MAS with glutamate/malate provoked an even stronger response. Malate, which on its own is unable to feed the ETC (*29*), and oxamate, which is not metabolized, both induced acidification of the mitochondrial matrix, consistent with H^+^-coupled entry (fig. S17). As expected, the alkalinization was partially reverted by activation of the F_1_F_o_-ATPase with ADP and inorganic phosphate, equivalent to State 4 respiration (Fig. 4E). These results confirm that the pH change induced by substrates reflects the pumping of protons by the ETC. Noteworthy, the entry of protons via the MPC, one per pyruvate, is negligible compared to the 30 protons that are extruded by the ETC for every pyruvate that is metabolized to CO_2_ (*29*). Due to space limitations, this methodology will be described in detail elsewhere. Next, we tested the ability of lactate to drive the ETC. Indeed, lactate was able to alkalinize the matrix, albeit only in a fraction of the experiments and at a lower rate than observed with pyruvate or glutamate/malate (Fig. 4F). Critically, in energized organelles, lactate was without effect or, when applied at higher concentrations, it induced matrix acidification (Fig. 4G). Neuronal mitochondria also showed a robust ETC-mediated matrix alkalinization, but could not be energized by lactate (Fig. 4H-I; fig. S18). To investigate the possibility of lactate consumption by an independent approach, the steady-state was interrupted with UK-5099, as previously done to image pyruvate metabolism (fig. S19; *30*). As illustrated in Figures 5B, left panel, and fig. S20, this inhibitor-stop method confirmed that mitochondria can consume lactate.

**Fig. 4.**
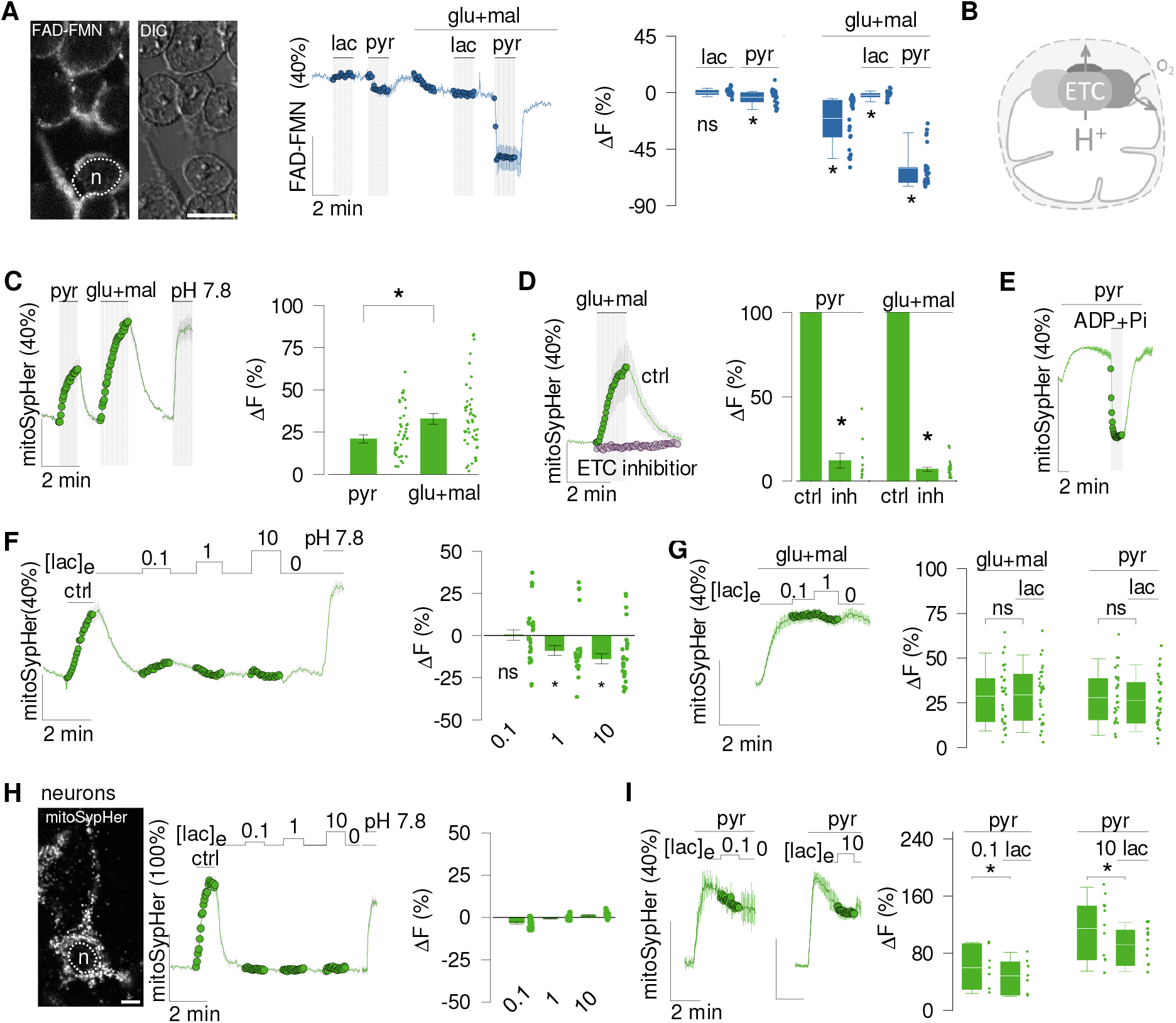
Weak mitochondrial energization by lactate. HEK293 cells were permeabilized with digitonin. (**A**) Left panel, image of FAD-FMN autofluorescence. A nucleus is indicated (n). Right panel, DIC image. Bar represents 20 µm. The graph shows the effect of pyruvate (0.1 mM) and lactate (0.1 mM) on FAD-FMN autofluorescence in the absence or presence of glutamate (0.2mM) and malate (0.1 mM). Summary of five experiments (28 cells; paired t-test). (**B**) Proton pumping by respiratory complexes is fueled by NADH and FADH_2_. (**C**) The effect of pyruvate (0.1 mM), glutamate (0.2 mM) and malate (0.1 mM), or buffer adjusted at pH 7.8 was measured with mitoSypHer. Calibrated signals obtained at pH 7.2 and 7.8 are indicated. Bar graph shows the changes after 90 seconds of substrate exposure (nine experiments, 47 cells). (**D**) Mitochondria were exposed to glutamate and malate in the absence or presence of ETC inhibitors: rotenone (1 µM) and antimycin (1 µM). Bar graph summarizes data for pyruvate (three experiment, 9 cells) and glutamate plus malate (five experiments, 20 cells), paired t-test. (**E**) Pyruvate-energized mitochondria were exposed to ADP (2 mM) and Pi (2 mM) (three experiments, 21 cells). (**F**) After a control pulse of glutamate and malate, mitochondria were exposed to increasing concentrations of lactate (mM). Bar graph shows the change in pH signal relative to baseline. Data from six experiments (28 cells), paired-t test. **(G)** Glutamate plus malate -energized mitochondria were exposed to lactate (mM). Bar graph shows the change in pH signal elicited by 0.1 mM lactate relative to baseline (five experiments, 28 cells, paired t-test). NS, not significant; *; p < 0.05. **(H)** Neurons expressing the pH sensor mitoSypHer. The nucleus is indicated (n). Bar represents 20 µm. After a control pulse of pyruvate, neurons were exposed to increasing concentrations of lactate (mM). (**I**)Bar graph summarized data from five experiments (21 cells, paired t-test).NS, not significant; *; p < 0.05. Bar graph shows the change in pH signal elicited by 0.1 mM lactate relative to baseline (five experiments, 28 cells, paired t-test). NS, not significant; *; p < 0.05.

**Fig. 5.**
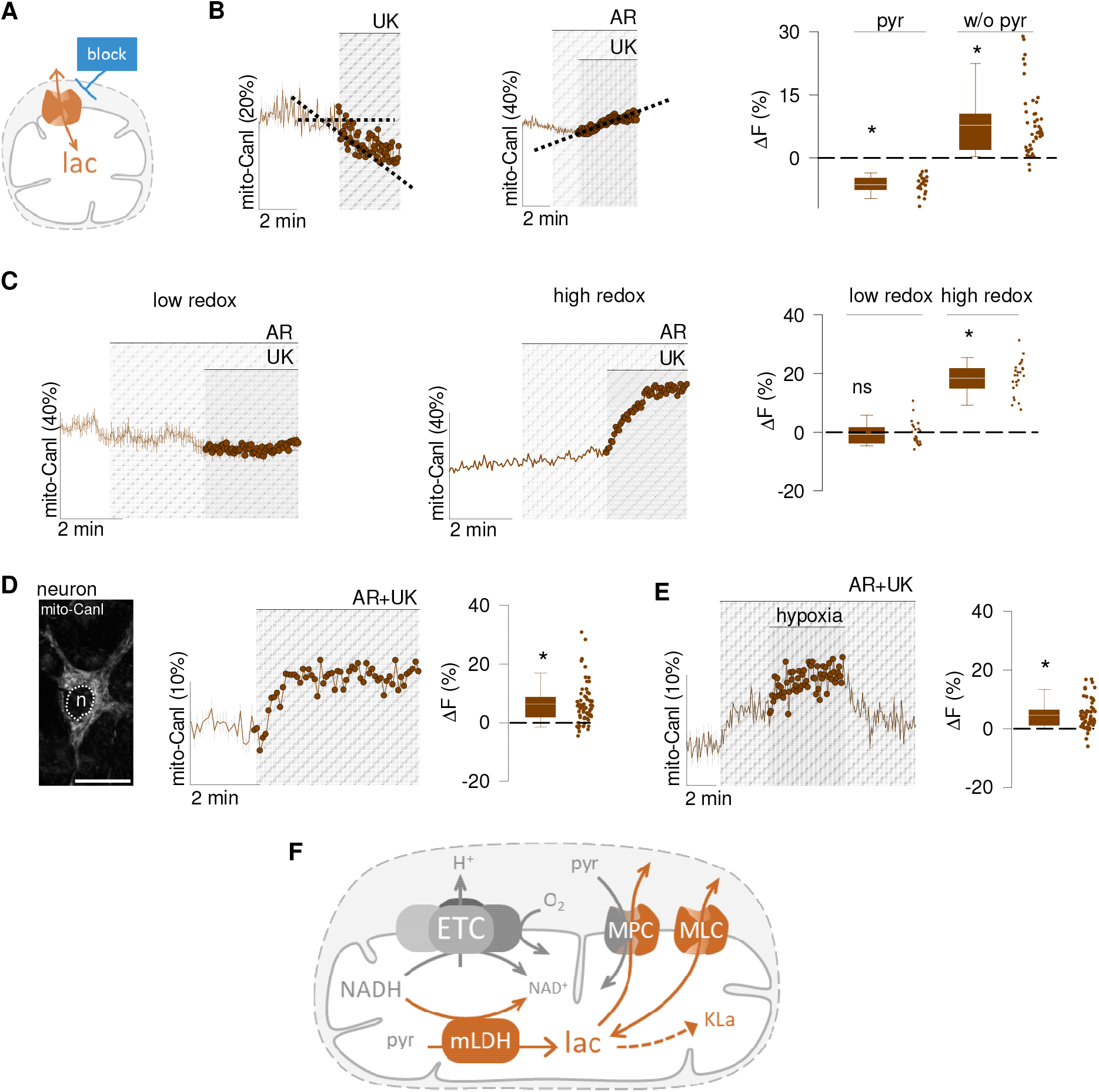
Lactate-carried mitochondrial overflow. (**A**) Net mitochondrial lactate flux in the presence of 0.5 mM lactate was determined in permeabilized cells expressing mito-CanlonicSF using an inhibitor-stop protocol (1 µM UK5099 and/or 1 µM AR-C155858). (**B**) Effect of lactate transport inhibitors on the steady-state of mitochondrial lactate in HEK293 cells in the absence (left panel) or presence (right panel) of 0.1 mM pyruvate. Perfusate contained 0.2 mM glutamate and 0.1 mM malate. Summary of data from five experiments without pyruvate (22 cells) and seven experiments with pyruvate (27 cells). Wilcoxon Signed Rank test; *, p < 0.05. (**C**) Effect of inhibitors on the steady-state of HEK293 mitochondrial lactate in the absence of glutamate/malate (low redox, left panel) or presence of glutamate/malate (high redox, right panel). Perfusate contained 10 mM pyruvate. Summary of data from three experiments at low redox (23 cells) and four experiments at high redox (24 cells). Paired t-test; *, p < 0.05. (**D**) Effect of transport inhibitors on permeabiized neurons expressing mito-CanlonicSF at high redox (glutamate/malate) plus 10 mM pyruvate. Bar represents 20 µm. Bar graph summarizes nine experiments (65 cells; paired t-test; *, p < 0.05). (**E**) Effect of hypoxia (N_2_-gassing) on the accumulation of lactate induced by transport inhibitors. Bar graph summarizes data from four experiments (34 cells, paired t-test; *; p < 0.05). (**F**) NADH fuels the electron transport chain (ETC) and also drives lactate production at mitochondrial LDH (mLDH). Lactate is then discarded via MPC and MLC. Protein lactylation (Kla) is determined by lactate level.

### Mitochondria are lactate producers: stimulation by hypoxia

When the transport of lactate was interrupted in mitochondria fueled by pyruvate, matrix lactate went up (Fig. 5B, right panel), reflecting direct-mode LDH, i.e. pyruvate to lactate conversion. In line with their relative strength at inhibiting mitochondrial lactate transport (Fig. 3F), UK-5099 was much more effective than AR-C155858. Note that because MPC blockage by UK-5099 is partial, there is enough pyruvate left in the matrix (fig. S19; *30*) to sustain PDH, which is a high affinity enzyme. Next we investigated the factors determining lactate production. In the absence of glutamate/malate, i.e. low matrix redox potential (see Fig. 4A), lactate was not produced (Fig. 5C, left panel). Strikingly, a reduced matrix led to robust lactate production (Fig. 5C, right panel). Similar experiments in neurons showed that their mitochondria are also net lactate producers (Fig. 5D). The permissive role of glutamate/malate, which provides electrons but not carbons, suggested to us that lactate serves to discard electrons, a kind of redox/energy overflow. This hypothesis was put to the test by exposing neuronal mitochondria to a solution bubbled with nitrogen, which in our perfusion system lowered the concentration of oxygen from 230 µM to 62 µM (*31, 32*). By impairing the acceptance of electrons at Complex IV, hypoxia jams the ETC, leading to accumulation of NADH. Consistent with the overflow hypothesis, hypoxia reversibly stimulated the production of lactate by mitochondria (Fig. 5E; fig. S21), but did not affect the pH of the organelle (fig. S22). A summary of the main conclusions of this work is presented in Fig. 5F.

## DISCUSSION

We looked for lactate consumption in mitochondria, but found that they are lactate producers. This conclusion is based on the following observations. i. A dynamic lactate pool in the matrix. ii. A high-affinity lactate transport system in the inner membrane. iii. MPC-sensitive lactate production from pyruvate. iv. Redox-sensitive lactate production from pyruvate. v. Stimulation of lactate production by hypoxia. In addition, we observed that lactate induces the lactylation of mitochondrial proteins, and that neuronal mitochondria sense blood lactate levels. The intracellular lactate shuttle (ILS) hypothesis states that lactate is transported into the mitochondrial matrix and then metabolized to pyruvate. Evidence supporting ILS include immunohistochemical detection of MCTs and LDH, and respirometry of mitochondrial preparations (reviewed by *10*). Issues regarding ILS include antibody specificity and purity of mitochondrial preparations (*4, 33*), whereas isotopic studies have given conflicting results (*8, 34*). A revised version of ILS has LDH in the intermembrane space (*10, 35*). The current work may help to reconcile some of these conflicting results.

### Lactate and pyruvate transport

Mitochondrial lactate was found to be well connected with the cytosol, just as well as cytosolic lactate is connected with the extracellular space. The saturable nature, affinity in the low millimolar range, and ability to carry protons, pyruvate and oxamate, first pointed to a monocarboxylate transporter. Previously, antibodies against MCT1 and MCT2 had detected epitopes associated with mitochondria (*9, 36*) and lower mitochondrial lactate was reported in skeletal muscle genetically-depleted of MCT1 (*37*). However, we do not favor MCT involvement. At 0.8 mM, the K_M_ of lactate uptake lies in the range of MCT2 (0.5-0.75 mM) and MCT4 (0.7-1 mM), and is lower than that reported for MCT1 (3.5-10 mM; *19, 22*). The robust mitochondrial uptake observed in MDA-MB-231 cells, which do not have the MCT1gene (*19, 23*), speaks against a mandatory role for this isoform, and so do the weak inhibitory effects of the MCT1/2 inhibitors AR-C155858 and AZD3965. Syrosingopine and pCMBS, which block MCT1 and MCT4, but not MCT2, also achieved partial inhibition. The best effect was achieved with the MPC blocker UK-5099, whereas genetic deletion of one of its subunits reduced lactate uptake by about 50%. These results suggest that there is significant lactate permeation via the MPC. Because of the standing inward pyruvate gradient (*38*) and the effective trans-acceleration of lactate by pyruvate, we surmise that the MPC is better poised to work as a lactate extruder (Fig. 5F). The remaining lactate permeability, which does not appear to be MCT1, MCT2 or MCT4, could tentatively be termed Mitochondrial Lactate Carrier (MLC; Fig. 5F).

### Roles of mitochondrial lactate

We argue here that the functional module consisting of matrix LDH and MLC plays the role of de-energizing mitochondria. Recently, the pyruvate concentration and NADH/NAD^+^ ratio in the matrix have been determined, which enables a thermodynamic analysis. The net direction of the LDH reaction is governed by its equilibrium constant and the concentrations of pyruvate, lactate, NADH, NAD^+^ and H^+^. With the equilibrium constant (K_eq_) of 1.11 × 10^−11^ M and a NADH/NAD^+^ ratio of 0.125 (*39*), pH at 7.8 (*40*), and matrix pyruvate at 30 µM (*30*), mitochondria should be able to push lactate up to 68 mM, almost two orders of magnitude higher than that observed by us and others (*41*). This means that mitochondrial LDH works far from equilibrium, which explains the modest or absent energization observed when the enzyme is forced to run in the reverse mode by applying lactate in the absence of pyruvate, and a more favorable NADH/NAD^+^ ratio of 0.009 (*42*). Low mitochondrial LDH activity also agrees with the previous difficulty on isolating the enzyme against the background of the very abundant cytosolic LDH, which works near equilibrium. Our results show that under substrate conditions prevailing in healthy cells, mitochondrial LDH runs in the direct mode, slowly converting NADH into NAD^+^, so that electrons are passed to lactate, which then leaves the matrix and the cell. What could be the point of such continuous waste of energy and carbons? A key to this question is suggested by the observed effect of hypoxia. When oxygen is low, electrons accumulate as NADH. Cells are known to reduce such redox stress by expressing uncoupling proteins (UCPs) that shortcut the ETC, bringing down matrix NADH levels and the production of reactive oxygen species (*32*). To exert their protective role, UCPs use oxygen and dissipate the energy irreversibly, as heat. The lactate overflow is faster, does not require precious oxygen, and saves energy as a transient lactate pool that may be shifted within the tissue and beyond. LDH is highly regulated and its partial pharmacological inhibition has consequences (*43, 44*), observations that have been puzzling because neither regulation nor partial inhibition should affect flux in a near-equilibrium reaction. In contrast, mitochondrial LDH works far from equilibrium. Now that a functional assay specific for mitochondrial LDH is available, it will be easier to investigate its biology and regulation, for example by lactylation. Lactate can stimulate the ETC directly, acting as a signal independently of its metabolism (*8*). The ability of mitochondria to generate their own lactate suggests a possible mechanism for the control of respiration and/or free radical production in subcellular domains.

## Acknowledgements

We thank members of the Energy Metabolism Group at CECs for helpful discussions, and Karen Everett for critical reading of the manuscript. This work was funded partly by Fondecyt projects 1230145 (LFB), 11190678 (IR) and 1230682 (IR), NIH1R01NS126920-01A1 (Phil O’Herron, PI; LFB, co-I), Proyecto USS-FIN-23-FAPE-03 (PYS). We also thank Fondequip EQM190142 (CAN).

## List of Supplementary Materials

Fig. S1. Colocalization of mito-CanlonicSF with a mitochondrial marker.

Fig. S2. Lactate pool quantification I.

Fig. S3. Lactate pool quantification II.

Fig. S4. Kinetics of mitochondrial lactate uptake.

Fig. S5. No detectable effect of lactate exposure on cytosolic and mitochondrial Ca+2

Fig. S6. Quantification of lactate-dependent whole-cell HEK293 protein lactylation.

Fig. S7. Proteomic identification of lactylated mitochondrial proteins.

Fig. S8i. Protein sequence coverage of human Peptidyl-prolyl cis-trans isomerase (FKBP4).

Fig. S8ii. Protein sequence coverage of human Stress-70 protein, mitochondrial (GRP75).

Fig. S8iii. Protein sequence coverage of human mitochondrial 10 kDa heat shock protein (CH10).

Fig. S8iv. Protein sequence coverage of human mitochondrial 60 kDa heat shock protein (CH60).

Fig. S8v. Protein sequence coverage of human L-lactate dehydrogenase B chain (LDHB).

Fig. S8vi. Protein sequence coverage of human mitochondrial Leucine-rich PPR motif-containing protein (LPPRC).

Fig. S8vii. Protein sequence coverage of human superoxide dismutase [Cu-Zn] (SODC).

Fig. S8viii. Protein sequence coverage of human small ribosomal subunit protein uS3 (RS3).

Fig. S8ix. Protein sequence coverage of human heat shock protein HSP 90-alpha (HS90A).

Fig. S8x. Protein sequence coverage of human mitochondrial malate dehydrogenase (MDHM).

Fig. S9. Digitonin permeabilizes the plasma membrane without apparent damage to mitochondria.

Fig. S10. Estimation of the lactate transport rate in energized mitochondria is reproducible.

Fig. S11. Validation of lactate transport blockers in intact cells.

Fig. S12. Partial inhibition of lactate transport in permeabilized cells.

Fig. S13. Differential effect of syrosingopine on permeabilized and intact MDA-MB-231 cells.

Fig. S14. MPC2KO inhibits the initial rate of pyruvate uptake.

Fig. S15. MPC2KO decreases the steady-state of lactate.

Fig. S16. Stability of mitochondrial energization by pyruvate.

Fig. S17. Effect of oxamate and malate on matrix pH.

Fig. S18. Effect of ETC inhibitors in energized neurons.

Fig. S19. Mitochondrial pyruvate consumption.

Fig. S20. No effect of DMSO on the lactate steady-state.

Fig. S21. Transient accumulation of lactate by hypoxia.

Fig. S22. Hypoxia does not affect matrix pH.

