## Supplementary material for "Lactate-carried Mitochondrial Energy Overflow": Materials and Methods and Supplementary figures

### Materials

Most reagents were purchased from Sigma-Aldrich/MERCK. Sodium L-lactate (Sigma-Aldrich (CAS N° 867-56-1), Sodium pyruvate (Sigma-Aldrich CAS N° 113-24-6), Sodium oxamate (Sigma-Aldrich CAS N° 565-73-1), L-glutamic acid monosodium salt monohydrate (Sigma-Aldrich CAS N° 6106-04-3), L-(-)-malic acid (Sigma-Aldrich CAS N° 97-67-6), and Sodium 3-hydroxybutyrate (Sigma-Aldrich CAS N° 150-83-4) were diluted in nano-pure water to make 1 M stocks. UK-5099 (Sigma-Aldrich CAS N° 56396-35-1), AR-C155858 (Sigma-Aldrich CAS N° 496791-37-8), AR-C141990 hydrochloride (Tocris Bioscience Cat N° 5658), AZD3965 (MedchemExpress Cat N° HY-12750), 4-(Chloromercuri) benzenesulfonic acid sodium salt (pCMBS; Toronto Research Chemicals Cat N° 14110-97-5), syrosingopine (Sigma-Aldrich CAS N° 84-36-6), rotenone (Sigma-Aldrich CAS N°83-79-4) antimycin A (Sigma-Aldrich CAS N°1397-94-0), adenosine 5' diphosphate sodium salt (Sigma-Aldrich CAS N°20398-34-9), ATP magnesium salt (CAS N°74804-12-9), FCCP (Sigma-Aldrich CAS N°370-86-5) and digitonin (Sigma-Aldrich CAS N°11024-24-1), were dissolved in DMSO to make 1000x stocks. Routine control experiments were carried out to rule out confounding effects of DMSO.

### Cell cultures

Human Embryonic Kidney cells (HEK293), MDA-MB-231, Cos-7, and Hep-G2 cells were purchased from the American Type Culture Collection (ATCC). Cultures were passaged with 1% trypsin at 37°C, followed by mechanical disaggregation. HEK293 were harvested in DMEM/F12 low glucose plus 10% fetal bovine serum (FBS). MDA-MB-231, Cos-7, and Hep-G2 cells were cultured in Leibovitz medium, high glucose DMEM and EMEM, respectively. Coverslips were ethanol sterilized and coated with 0.01% poli-L-lysine. Culture were grown at 37°C in 95% air, 5% $CO_2$, except for MDA-MB-231, which were cultured in 100% air. Experiments were carried out at 40%-60% confluence

### Reporter cell lines and MPC2KO

Stable cell lines were derived from HEK293 cells by lentiviral bicistronic-vector infection (*1*). Cells were subcultured to obtain a final confluence of 40-60%. HEK040 expresses mito-CanlonicSF, HEK024 expresses CanlonicSF (Addgene plasmid #178342) in the cytosol (*2*), HEK017 expresses Pyronic (Addgene plasmid #51308) in mitochondria, and MDA021 expresses PyronicSF (Addgene plasmid #124812) in the cytosol. Genetic deletion of the MPC2 was induced by CRISPR/Cas9 edition followed by puromycin (2 ug/ml) selection. The plasmids used for the genetic edition of MPC2 were gifted by Jason Cantor (*3*): pLentiCRISPR-v1-sgMPC2_7 (Addgene plasmid # 163457) and pLentiCRISPR-v1-sgMPC2_9 (Addgene plasmid # 163458).

### Neural cultures

Animal handling was carried out in strict accordance with the recommendations in the Guide for the Care and Use of Laboratory Animals of the National Institutes of Health. Procedures were approved by the Centro de Estudios Cientıficos Animal Care and Use Committee, project 1230145, and sanctioned by the Dirección de Integridad, Seguridad y Etica de la Investigación, Universidad San Sebastián.To obtain cultures enriched in neurons, hippocampus or cortical tissue was extracted from F1 crossover C57BL/6J × CBA/J embryos (18 days). The tissue was dissected from meninges, maintained in an ice-HBSS medium, and enzymatically dissociated with 1% trypsin.

After mechanical disaggregation, cells were cultivated in a glass coverslip with neurobasal medium plus 2% B27 supplement, 10 mM glucose, 1% glutamax, and antibiotics (2.5 mg/ml fungizone and 10 mg/penicillin/streptomycin). The medium was changed every three days, and cells were used after day twelve of cultivation at 95% air and 5% $CO_2$. For astrocyte enrichment on neuron cultures, the media was supplemented with 3% FBS. To obtain cultures enriched in astrocytes, cortical tissue was extracted from eighteen-day-old embryos (F1 crossover C57BL/6J × CBA/J). The cortex was separated from the meninges, hippocampus, and medulla oblongata and maintained in an ice-HBSS medium for enzymatic dissociation. Cells were grown on coverslips with neurobasal medium plus 10% FBS, 2% B27 supplement, 10 mM glucose, 1% glutamax, and antibiotics (2.5 mg/ml fungizone and 10 mg/ml penicillin/streptomycin). The medium was changed every three days, and cells were used after day eight of cultivation at 95 % air and 5% $CO_2$.

<u>Transfection/infection for in vitro sensor expression</u>
Reagents were acquired from Invitrogen. The mix reaction included the plasmid: MitoSypHer (2 μg; Addgene # 48250), lipofectamine 3000 (2 μL), P3000 reagent (2 μL) and Opti-MEM medium (400 μL). Transfection was done at low confluence (25%), followed by 16-24 hour incubation. Mito-CanlonicSF baculoviral particles were generated in the laboratory. Cell lines were incubated for 48 hours before experiments. Astrocyte cultures were infected between the sixth- and seventh-day for 48 hours with no medium change, and primary neuron cultures were infected between the eighth- and ninth-day for 72 hours without changing the neurobasal medium.

<u>In vivo assays</u>
All in vivo experimental and surgical procedures were approved by the local veterinary authorities in Zurich according to the guidelines of the Swiss Animal Protection Law, Veterinary Office, Canton of Zurich (Animal Welfare Act 16 December 2005, and Animal Welfare Ordinance 23 April 2008). Female C57BL/6J mice (Charles River) of 8-16 weeks of age (20-27 g bodyweight) were used in this study. All animals were kept in standardized IVC cages with access to water and food at libitum and were subjected to an inverted 12 h light-dark cycle.

Anesthesia. For both surgical interventions and in vivo 2-photon imaging, animals were anesthetized with a mixture of fentanyl (0.05 mg/kg bodyweight; Sintenyl, Sintetica), midazolam (5 mg/kg bodyweight; Dormicum, Roche), and medetomidine (0.5 mg/kg bodyweight; Domitor, Orion Pharma), which was injected subcutaneously (s.c.). Vitamin A ointment (VitA POS, Pharma Medica) was applied to both eyes. Oxygen was supplied throughout anesthesia and animals were kept on a homeothermic blanket until fully recovered.

Headplate implantation, craniotomy, and virus injection. To allow reproducible conditions for in vivo imaging, prior to the craniotomy, a custom-made aluminum head plate was implanted. Mice were fixed in a stereotactic frame (Model 900; David Kopf Instruments). After fur removal, the skin of the scalp was disinfected (Kodan; Schülke & Mayr) and local anesthesia was applied prior to midline incision. The exposed skull was cleaned, and a blue light curable bonding agent applied (Gluma Comfort; Heraeus Kulzer). Next, the head-plate was attached using light-curing dental cement (Tetric EvoFlow; Ivoclar Vivadent). Above the left somatosensory cortex, a craniotomy was performed using a dental drill (OSSEODOC; Bien-Air). Adeno-associated viral vectors

(AAVs) were injected intracortically at 350 µm and 150 µm below the dura (80 nl each) by a custom-made micro injector to achieve neuronal sensor protein expression.

The following constructs were used: AAV6/2-hSyn1-chl-Mito-CanlonicSF-WPRE-bGHp (titer: 2.55 x 1012 vg/ml), and (in a subset of mice) additionally AAV1/2-hSyn1-Laconic-WPRE-hGHp (titer: 3.55 x 1012 vg/ml). After virus injections, the brain was covered by a square sapphire glass (3 x 3 mm; 19395-1, Hebo Special Glass), which was sealed with dental cement. In mice used for histology, no headplate was implanted and viral constructs were intracortically injected via 3 small drill holes. For postoperative analgesia, mice were injected s.c. with buprenorphine (0.1 mg/kg bodyweight; Temgesic, Indivior Schweiz AG) and Carprofen (10 mg/kg bodyweight; Rimadyl inj. ad us. vet., Pfizer) directly after surgery and with Carprofen every 12 hours thereafter until fully recovered.

Experimental protocol. For imaging experiments and blood plasma lactate measurements after a baseline of 1 min, a 375 mM sodium L-lactate (L7022, Sigma-Aldrich) solution was injected at 1.5 mmol/kg bodyweight over 3 min intravenously via a tail vein catheter using a manually operated peristaltic pump (Reglo digital ISM831, Ismatec SA). For experiments in which plasma lactate levels were determined, the femoral artery was exposed and cannulated with fine bore polyethylene tubing (0.28 mm ID, 0.61 mm OD; Portex; Smiths Medical). Drops of blood were collected from the arterial catheter and blood plasma lactate was measured using an enzymatic lactate assay (Lactate pro-2, Arkray Healthcare, USA) at varying time points during the above-mentioned protocol.

Cerebral in vivo lactate measurements. Mice were imaged 3-4 weeks after virus injection using a custom-built 2-photon laser scanning microscope (*4*) with a tunable pulsed laser (Chameleon Discovery NX TPC; Coherent) at 920 nm excitation wavelength and equipped with a 16x (N16XLWD-PF, 0.8 NA, Nikon) or 25x water immersion objective (W-Plan-Apochromat 25x/1.05 NA, Olympus). Excitation and emission beam paths were separated by a dichroic mirror (F73-825; AHF Analysentechnik). Emission was further separated by dichroic mirrors at 506 nm (F38-506; AHF Analysentechnik) and at 560 nm (F38-560; AHF Analysentechnik) and was detected with photomultipliers (H9305-03, Hamamatsu) equipped with respective emission filters for blue (F39-477; AHF Analysentechnik), and for green (F37-545; AHF Analysentechnik). During imaging, mice were head-fixed and kept under anesthesia as described above. Anatomical images were acquired at 0.74 Hz and 512 x 512-pixel resolution. Time series for Canlonic measurements alone were acquired at 1.48 Hz (256 x 256-pixel resolution). Experiments comparing Canlonic with Laconic signal changes were acquired at 0.53 Hz (256 x 256-pixel resolution) cycling between the two sensor expression areas. All experiments were acquired using ScanImage (r3.8.1; Janelia Research Campus; (*5*) for 10 min.

<u>Immunohistochemistry</u>
After three weeks of virus injections, mice were deeply anesthetized with 200 µl pentobarbital (50 mg/ml intraperitoneally; Kantonsapotheke Zürich) and transcardially perfused with artificial cerebrospinal fluid (ACSF, pH 7.4) followed by 2 % paraformaldehyde (PFA, dissolved in 1 X phosphate-buffered saline (PBS), pH 7.4). Following perfusion, the brain was carefully extracted, post-fixed in 4 % PFA for 3 h and subsequently kept in a 30 % sucrose solution (in 1 X PBS, pH 7.4) overnight at 4°C. The fixed tissue was then frozen and cut into 30 µm coronal sections using

a cryomicrotome (Hyrax KS 24 microtome; Zeiss, Switzerland). Immunohistochemistry was performed according to standard protocols for staining of fixed, free-floating sections. In short, sections were washed, pre-blocked for 1 h (using normal donkey serum), and then incubated with primary antibodies (see below) overnight at 4°C. The next day, sections were washed, incubated for 45 min with secondary antibodies and DAPI (abcam, 228549), washed again, mounted onto glass slides, and coverslipped with mounting medium (Dako fluorescence mounting medium; Agilent, CA, USA). Images were acquired using a confocal laser scanning microscope (LSM 800; Zeiss, Switzerland) equipped with a 63x objective (Plan-Apochromat, NA 1.4, Oil). Primary antibodies: rabbit anti-NeuN (abcam, 177487), goat anti-GFP (Thermo Fisher, 600-101-215M), and mouse anti-HSP60 (abcam, 59457). Secondary antibodies: Alexa Fluor 647 donkey anti-rabbit (Invitrogen, A32795), Alexa Fluor 488 AffiniPure donkey anti-goat (Jackson ImmunoResearch, 705-545-03), and Cy3 AffiniPure donkey anti-mouse (Jackson ImmunoResearch, 715-165-151).

Fluorescent dye loading

BCECF, calcein, FLUO-4, Rhod-2 and TMRM were acquired from Sigma-Aldrich. The dyes were ester loaded. Stocks were prepared in DMSO: BCECF (2 mM), calcein (4 mM), FLUO-4 (4 mM) and 40 mM (Rhod-2). These stocks were re-dissolved in pluronic F-127 20% within the corresponding saline buffer for 10 minutes in a blackout recipient at 37°C. The final concentration to obtain was 400 nM, and 400 µM for Rhod-2- The procedure includes a de-esterification step. Rhod-2 was diluted in KRH glucose 5mM, and cultures were incubated for 15 minutes at 4°C. The solution was removed, and the corresponding cellular medium was added overnight for 16 hours. TMRM (Tetramethylrhodamine methyl ester perchlorate) was diluted at a saline buffer at a final concentration of 40 nM, with an incubation period of 1-3 minutes. Imaging was taken every few seconds until reached maximal brightness intensity.

Cellular permeabilization

Cultures were washed with KRH buffer of the following composition (mM): 134 NaCl, 5 KCl, 1.25 MgSO4, 1.25 CaCl2, 10 HEPES at pH 7.40. For permeabilization cultures were perfused for 3 min with 30 µM digitonin and 200 µM ADP in intracellular solution (mM): 130 KCl, 10 NaCl, 1.25 $MgCl_2$, 0.37 $CaCl_2$, 1 EGTA, 10 HEPES at pH 7.20. Estimated free $Ca^{2+}$ in this solution is 90 nM (Maxchelator). Digitonin was washed out for at least 10 minutes in intracellular buffer. The osmolarity of the buffer was between 284 and 286 mOsm/L.

In vitro imaging

Experiments were performed in Olympus IX70 and FV1000 confocal microscope at 22-25°C. The Olympus IX70 is an inverted fluorescence microscope equipped with a mercury/xenon arc lamp, pairs of excitation/emission filters, and a 20X oil-immersed objective. The FV1000 confocal microscope has a scanner laser for excitation at 440/488/543 within an upright 20X water-immersed objective (N.A 1.0). Both microscopes have an integrated perfusion and drainage system to manipulate substrate and compound irrigation. Images were obtained using the pinhole at the maximal opening. Software settings were 488 (3%) and 543 (10%), digital zoom 4X, 320*320 (pixels), and imaging was taking every 3, 5 or 10 seconds.

Metabolite sensor imaging. Most of the sensors used were GFP-based; therefore, most proteins were excited at 488 nm. Mito-SyPher was excited at 488/440 nm sequential line, emission wavelength 510-525/505-510. The light emitted at 440 nm corresponds to the isosbestic point, that was used for ratiometric measurements. Mito-CanlonicSF was excited sequentially at 488/543 with emission at 505-525/560-660 nm. Mito-PyronicSF was imaged at 505-525/560-660 nm (excitation laser line 488/543). All sensors have a second line reader (red protein or isosbestic point). This measurement is used as a baseline to dismiss artifacts.

Fluorescent dye imaging. BCECF was excited at 488/440, emission 510-525/505-510. Calcein and FLUO-4 as green-fluorescent dyes were imaged at 510-525. TMRM and Rhod-2 were excited at 543 and imaged at 560-660 nm.

Autofluorescence imaging. Flavoprotein autofluorescence was measured at 505-525 nm under high laser power (10% of the 488 nm line).

<u>Calibration of mito-CanlonicSF</u>
A multiplate reader (Perkin Elmer) was used to determine mito-CanlonicSF $K_D$ in mitochondria. HEK040 were cultured in 96 well plates and permeabilized under glutamate 0.2 and malate 0.1 mM. First reads correspond to nominal zero (6 mM of oxamate) and second reads involved curves dose-response of lactate with or without 1 μM of FCCP. Lactate concentrations were (mM): 0.0375, 0.07, 0.1, 0.3, 0.6, 1.2, 2.4, 10, 20 and 100. Each well was excited at 488 and 543 nm . The ratio of the emission length was obtained, and the data was normalizing with the first reads. The average of each dose was calculated, processed in percentage (%). A rectangular hyperbola was fitted to the date to obtain the $K_D$.

<u>Lactylation analysis</u>
Western Blotting. HEK293 cells ($2x10^6$) were treated in the presence or absence of lactate. Cells were resuspended (1X PBS) in agitation and centrifuged at 850g for 2 minutes (2X). Mitochondrial fractioning was performed by density gradient centrifugation. The fraction was lysed in RIPA (1% SDS + 1% deoxycholate and 1% triton), re-suspended in 100 uL of RIPA modified-buffer and vortex for 1 min. The sample were centrifugate at maximal speed. Proteins were quantified by QuantiPro BCA Assay Kit (Sigma Aldrich, QPCA-1KT) and 15 μg of protein were loaded per well. The gel electrophoresis was carried out in polyacrylamide 6% and 12%. First antibody, Anti Kla rabbit 1:5000 (PTM Biolabs, catalog N° PTM-1401), was maintained in agitation overnight at 4°C. The second antibody, Horseradish peroxidase-conjugated 1:10000, was incubated at 37°C for 1 hour. West Femton Maximal Sensitivity substrate (peroxidase) (Thermo Fisher catalog N° 34095) was used for chemiluminescence visualization. β-actin (Santa Cruz biotechnology C4, catalog N° sc-47778) was blotted as an internal control. For Western blot quantification images were processed in Fiji. Each band was selected The selected area was kept identical to contrast reasonably each fraction and condition. The data obtained was the area under the curve normalized by β-actin.

Proteomics. HEK293 cells ($2x10^6$) were used for the control and lactate-treated samples. Cells were lysed in 8M Urea, 100 mM NaCl, 50 mM ammonium bicarbonate (ABC) and 1X protease inhibitor (complete Protease Inhibitor Cocktail, Roche). Sonication was performed by 10 cycles of 10 seconds each, one minute of rest. Centrifugation was done by 15 minutes of at 15,000g. The supernatants were collected and quantified using the Dual-Range™ BCA Protein Assay kit (ThermoFisher). Proteins (100μg) were reduced with dithiothreitol 20mM (30 min, 60°C) and alkylated with iodoacetamide 40 mM (15 min, room temperature in darkness). The samples were treated with urea (0,6 M) and trypsin in a ratio 1:50 with overnight agitation at 37°C. After 16h the reaction was stopped with trifluoroacetic acid (TFA). The digested proteins (200 µg) were used for immunoprecipitation using anti-Kla antibody coupled to ProtA/G-beads (Santa Cruz). Antibody was coupled to ProtA/G-beads overnight at 4°C in agitation and washed-out (3X) 24 hours later with ETN buffer (NaCl 3M, EDTA 0,1M and TRIS 1,5M, pH 8,0). The digested peptides were added and incubated for 6 hours at 4°C in agitation. Immunocomplexes were washed with ETN buffer (3X) and with ultrapure water (2X).The peptides were eluted using TFA 1% and cleaned-up using Pierce C18 Spin Columns (ThermoFisher).

For nHPLC-MS/MS analysis tryptic peptides (500 ng) were separated using an Easy nLC II liquid chromatograph and analyzed on a Q Exactive™ Plus mass spectrometer. Peptides were resolved using a C18 PepMap™ Easy-Spray reverse-phase column (75µm x15cm) with a particle size of 3µm in a gradient of 6-35% ACN over 50 minutes for proteome and 30 minutes for immunoprecipitation, followed by 35-45% over 15 minutes at a flow rate of 200nl/min for a total run of 98-minutes for the proteome and 78-minutes for immunoprecipitation analyses were used. The parameters were set using Xcalibur acquisition software (version 4.2). Spectra were obtained using the Orbitrap analyzer in FullMS mode within a range of 300-1800m/z at a resolution of 70,000. The 20 and the 5 most intense ions were selected for fragmentation in the proteome and the immunoprecipitation, respectively. dd-MS2 spectra were acquired at resolution of 17,500. The target value was set to 1x105 with an isolation window of 1.8m/z. The maximum injection time was set to 100 msec with a normalized collision energy of 28eV.

Database search. Spectra were searched against UniProtKB/Swiss-Prot database search restricted to Homo sapiens (released on 04/2022, with 20374 entries), using PEAKS Studio software (version 10.6) with a parent/fragment mass error tolerance of 10ppm/0.02Da. Potential false positives were filtered using a 1% False Discovery Rate (FDR). Variable proteome modifications included Oxidation (M), Pyro-glu form E, Pyro-glu form Q, Deamidation (NQ) and Lactylation (K) (+72.02 Da). The subcellular location of mitochondrial proteins was identified by using UniProtKB/Swiss-Prot database search.

Image Processing. Data is obtained from sequential images and managed in Fiji. The background or noise was substracted in each photo of every channel. Due to the low-resolution on which the light was captured to increase temporal resolution, a mitochondrial mask was applied. This mask allows thresholding the fluorescent signal for each channel in a specific pixel range to avoid outliers and dismiss noisy brilliance. The final outcome of the mask assigned a value of zero to those very low signals and was acquired as integrated density.

Statistical analysis
Data are expressed as mean ± SEM. Data analysis was performed in Sigma Plot software. A paired t-test was applied to ascertain differences in before-after treatment protocols with normal distribution. In case of failed normality test (Shapiro-Wilk), differences were assigned with the Mann Whitney-Wilcoxon signed rank test (pairs). *, $p < 0.05$. Not significant (NS), $p > 0.05$).

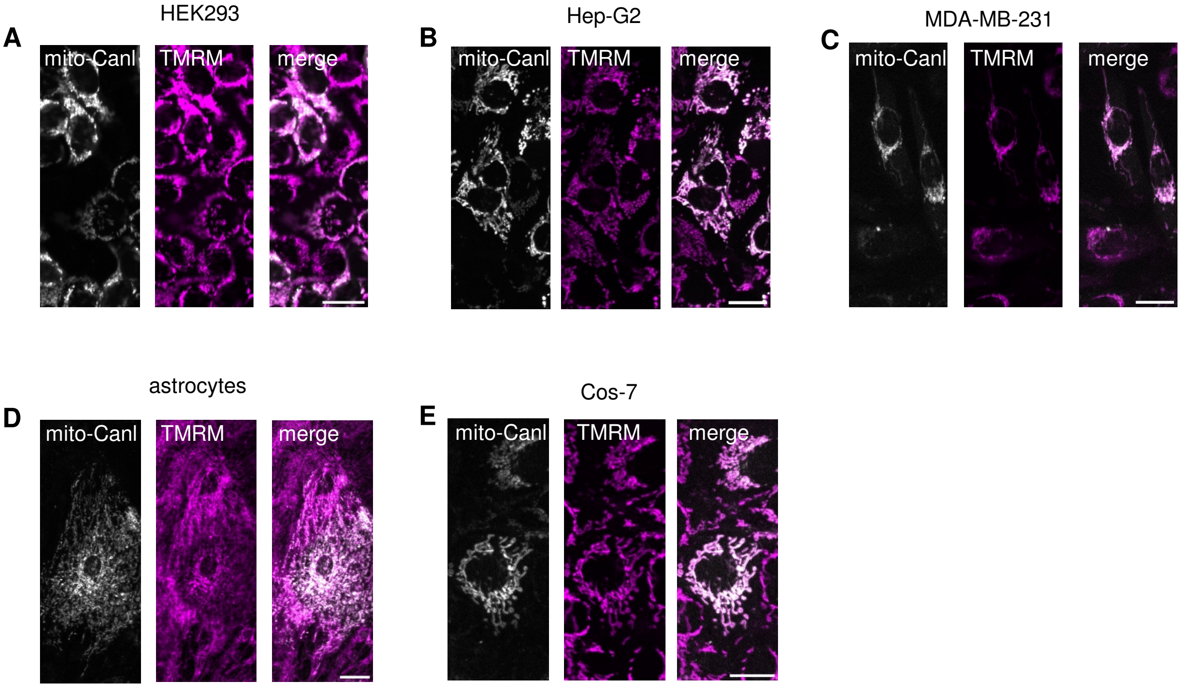


**Fig. S1. Colocalization of mito-CanIonicSF with a mitochondrial marker**. (**A**) HEK040 stable cell line expressing mito-CanIonicSF. (**B-E**) Mito-CanIonicSF was expressed with a baculovirus vector. Cultures were loaded with TMRM (see materials and methods). Bars represent 20 µm.

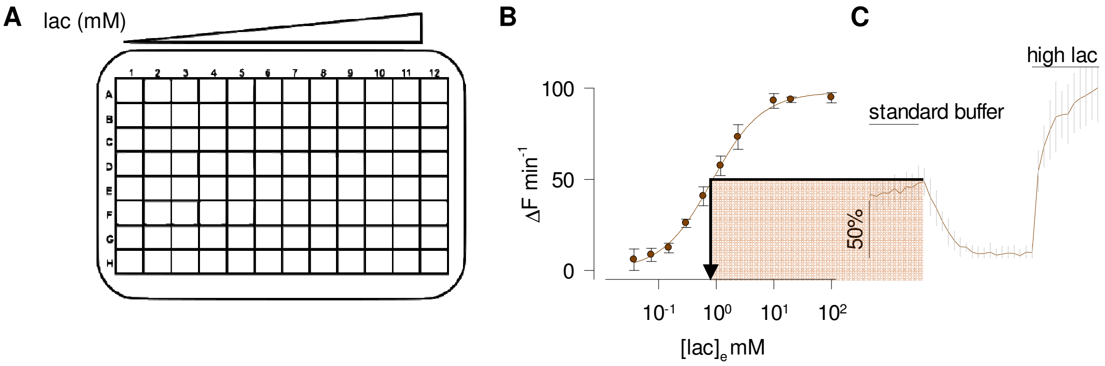


**Fig. S2**. **Lactate pool quantification I**. (**A**) Permeabilized HEK293 cells expressing mito-CanlonicSF were grown on 96 well-plates. After a first measurement in the presence of 6 mM oxamate (nominal zero lactate), cells were exposed to increasing lactate concentrations (mM) 0.0375, 0.075, 0.15, 0.3, 0.6, 1.2, 2.4, 10, 20, 100. (**B**) Normalized data from five experiments (mean ± SEM). Fitting of a rectangular hyperbola gave a $K_D$ of 1.4 ± 0.3 mM. (**C**) Example of lactate pool estimation. Astrocytes were sequentially exposed to standard buffer (2 mM glucose/0.05 mM pyruvate/0.5 mM lactate), 6 mM oxamate and 10 mM lactate (averaged trace from 3 cells). Interpolation of the fluorescence level in standard buffer gave a concentration of 0.8 mM (dotted line).

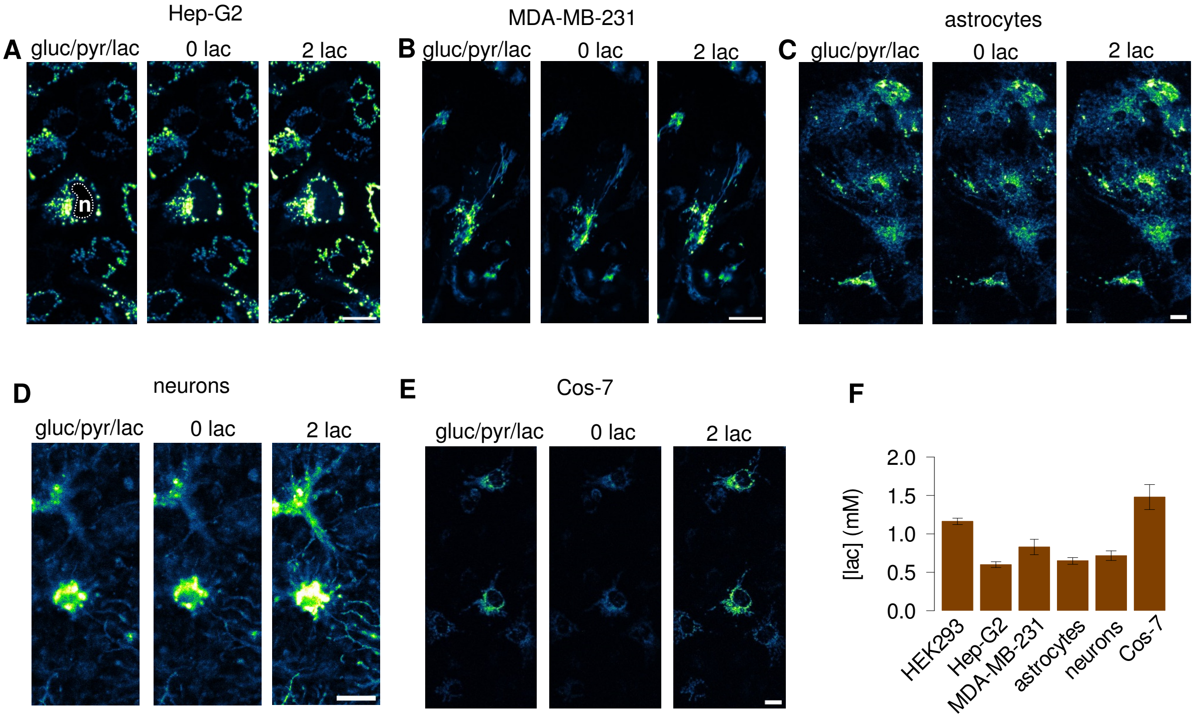


**Fig. S3. Lactate pool quantification II**. Intact cells expressing mito-CanlonicSF were sequentially imaged in the presence of standard buffer (5 mM glucose/0.05 mM pyruvate/0.5 mM lactate), 0 lactate and 2 mM lactate. Bar represents 20 µm. (**A**) Hep-G2 (three experiments, 51 cells). (**B**) MDA-MB-231 (three experiments, 26 cells). (**C**) Astrocytes (eleven experiments, 45 cells). (**D**) Neurons (eleven experiments, 19 cells). (**E**) Cos-7 (six experiments, 32 cells). In brain cells, standard buffer had 2 mM glucose/0.05 mM pyruvate/0.5 mM lactate. (**F**) Bar graph shows the estimated lactate pools.

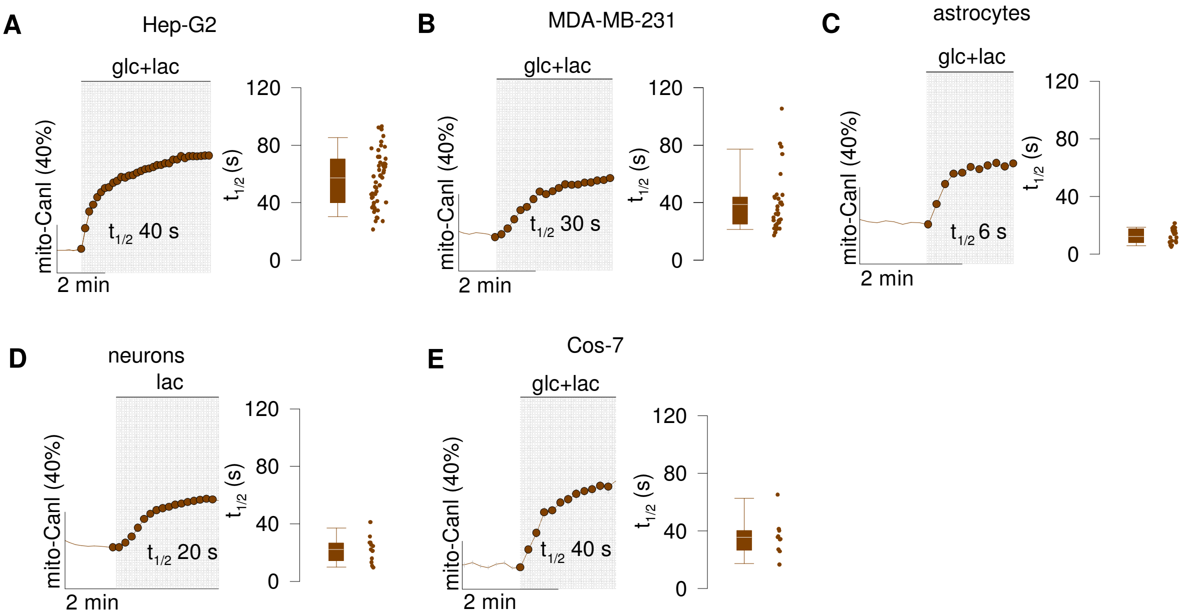


**Fig. S4. Kinetics of mitochondrial lactate uptake**. Intact cells were exposed to 5 mM glucose/5mM lactate or 2 mM lactate (brain cells). Representative traces and corresponding half-time. (**A**) Hep-G2 (three experiments, 51 cells). (**B**) MDA-MB-231 (three experiments, 32 cells). (**C**) astrocytes (four experiments, 24 cells). (**D**) Neurons (six experiments, 13 cells). (**E**) Cos-7 (two experiments, 13 cells).

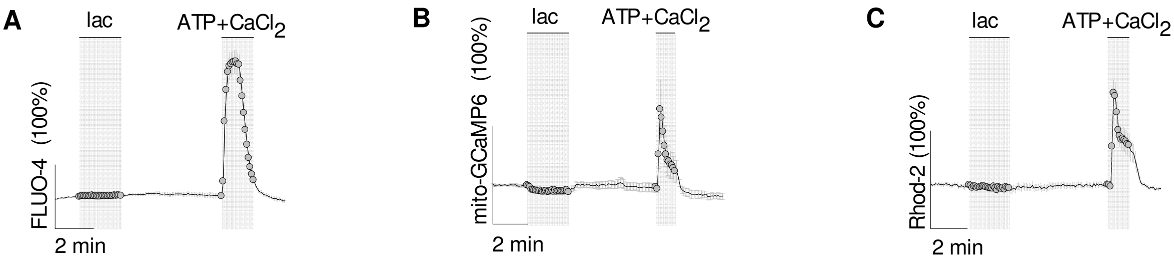


**Fig. S5. No detectable effect of lactate exposure on cytosolic and mitochondrial $Ca^{+2}$.** HEK293 cells were exposed to 1 mM lactate or 0.1 mM ATP plus 10 mM $CaCl_2$. **(A)** Cytosolic $Ca^{+2}$ was monitored with FLUO-4 (20 cells). **(B)** Mitochondrial $Ca^{+2}$ was monitored with mito-GCaMP6s (two experiments, 15 cells). **(C)** Mitochondrial $Ca^{+2}$ was monitored with Rhod-2 (three experiments, 32 cells).

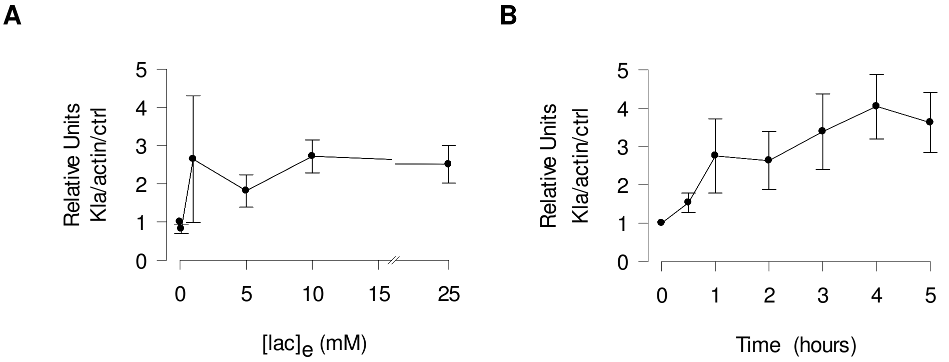


**Fig. S6. Quantification of lactate-dependent whole-cell HEK293 protein lactylation in response to lactate.** Signal was obtained by immunoblot densitometry. (**A**) Cells were exposed to 0.1, 1, 5, 10, and 25-mM lactate (mean ± SEM from three experiments). (**B**) Cells were exposed 25 mM lactate for the indicated times (mean ± SEM from seven experiments).

**A** Whole-cell protein-digest

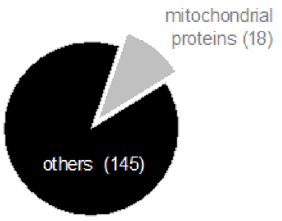


Kla immunoprecipitate

**B**

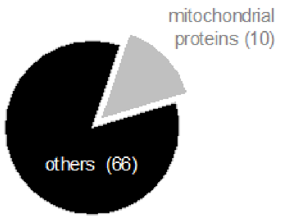


**C**

**Table 1**| Mitochondrial lactylated proteins

| Code | description | Kla position |
|---|---|---|
| Q02790 | Peptidyl-prolyl cis-trans isomerase (FKBP4) | 391 |
| P38646 | Stress-70 protein, mitochondrial (GRP75) | 646 |
| P61604 | 10 kDa heat shock protein, mitochondrial (CH10) | 36 |
| P10809 | 60 kDa heat shock protein, mitochondrial (CH60) | 72,75 |
| P07195 | L-lactate dehydrogenase B chain (LDHB) | 7,43 |
| P42704 | Leucine-rich PPR motif-containing protein, mitochondrial (LPPRC) | 431 |
| P00441 | Superoxide dismutase [Cu-Zn] (SODC) | 10 |
| P61247 | Small ribosomal subunit protein uS3 (RS3) | 195 |
| P08238 | Heat shock protein HSP 90-alpha (HS90A) | 107,203 |
| P40926 | Malate dehydrogenase, mitochondrial (MDHM) | 26,78 |

**Fig. S7. Proteomic identification of lactylated mitochondrial proteins.** HEK293 cells were treated with 25 mM lactate for five hours. Lactylated proteins were identified by nHPLC-MS/MS analysis of protein digests obtained by trypsination, as described in methods. (**A**) Whole-cell extracts were found to contain 163 lactylated proteins,18 of which were mitochondrial. (**B**) A subset of lactylated proteins were identified by immunoprecipitation of whole-cell protein digests using anti-lactyl-lysine antibody (Kla), of which 10 were mitochondria. (**C**) List of mitochondrial proteins from B. Protein accession numbers are from the UniProt database. Sites of lactylated lysines are indicated.

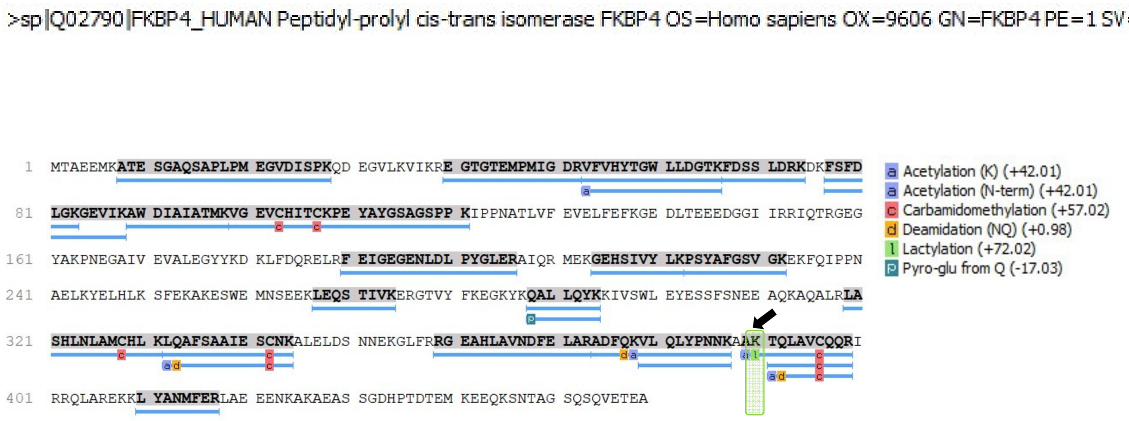


**Fig. S8i. Protein sequence coverage of human Peptidyl-prolyl cis-trans isomerase (FKBP4).** The amino acid sequence of human Peptidyl-prolyl cis-trans isomerase (FKBP4) is shown. Blue lines represent peptides obtained from a HEK293 protein digest. Post-translational modifications are indicated, were lactylation is highlighted.

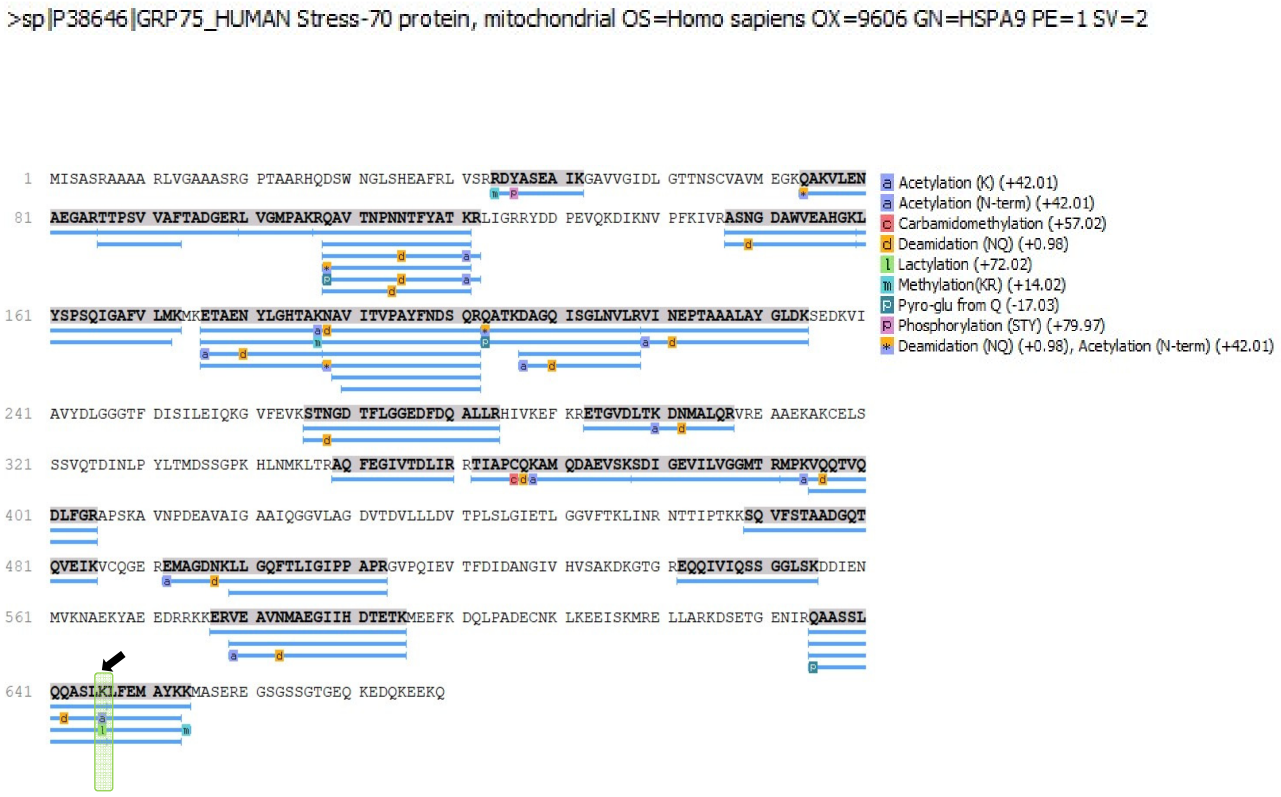


**Fig. S8ii. Protein sequence coverage of human Stress-70 protein, mitochondrial (GRP75).** The amino acid sequence of human GRP75. Blue lines represent peptides obtained from a HEK293 protein digest. Post-translational modifications are indicated, were lactylation is highlighted.

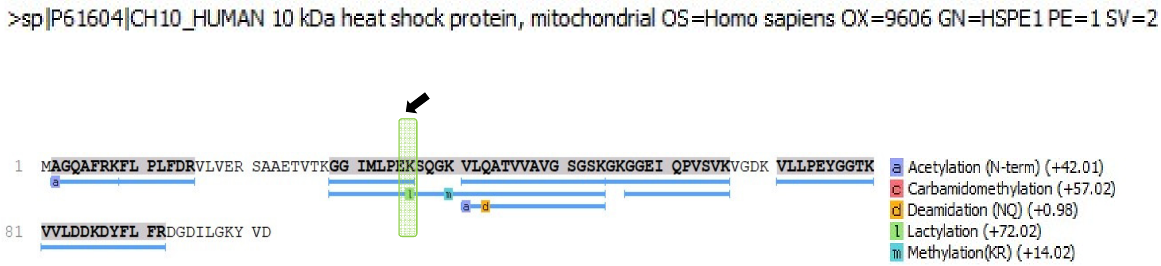


**Fig. S8iii. Protein sequence coverage of human mitochondrial 10 kDa heat shock protein (CH10).** The amino acid sequence of human CH10 is shown. Blue lines represent peptides obtained from a HEK293 protein digest. Post-translational modifications are indicated, were lactylation is highlighted.

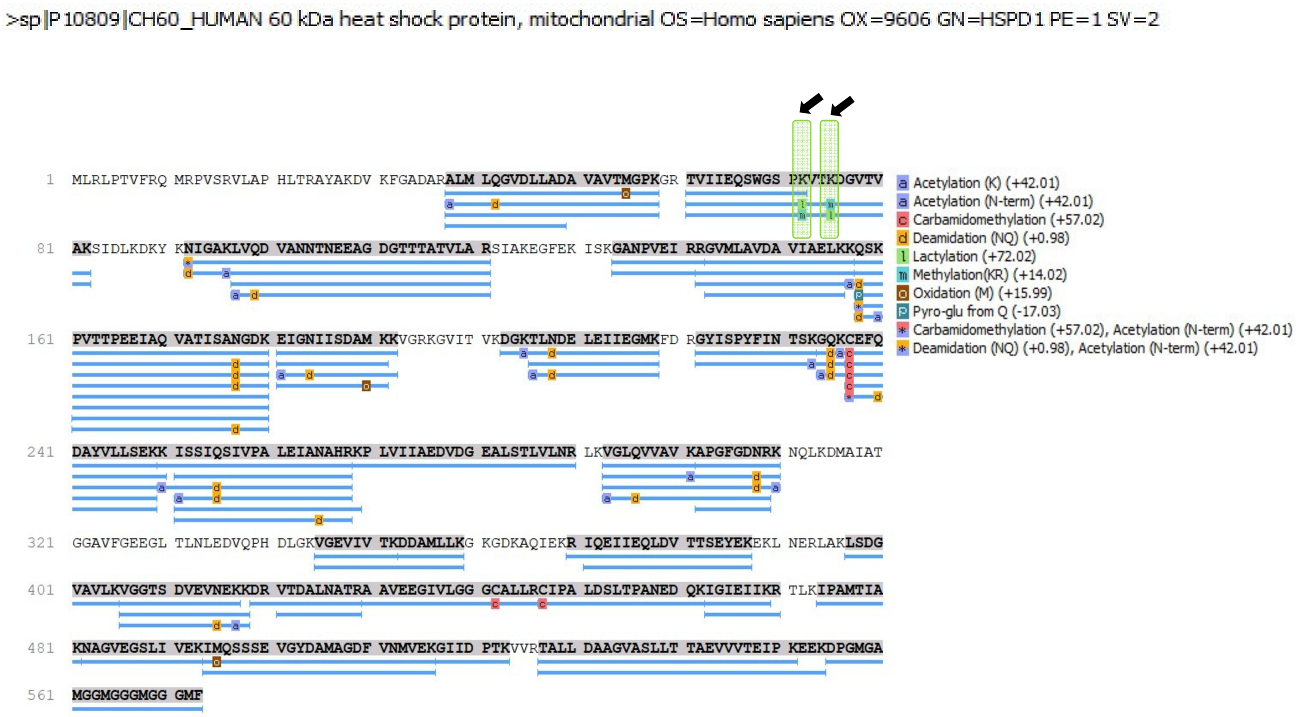


**Fig. S8iv. Protein sequence coverage of human mitochondrial 60 kDa heat shock protein (CH60).** The amino acid sequence of human CH60 is shown. Blue lines represent peptides obtained from a HEK293 protein digest. Post-translational modifications are indicated, were lactylation is highlighted.

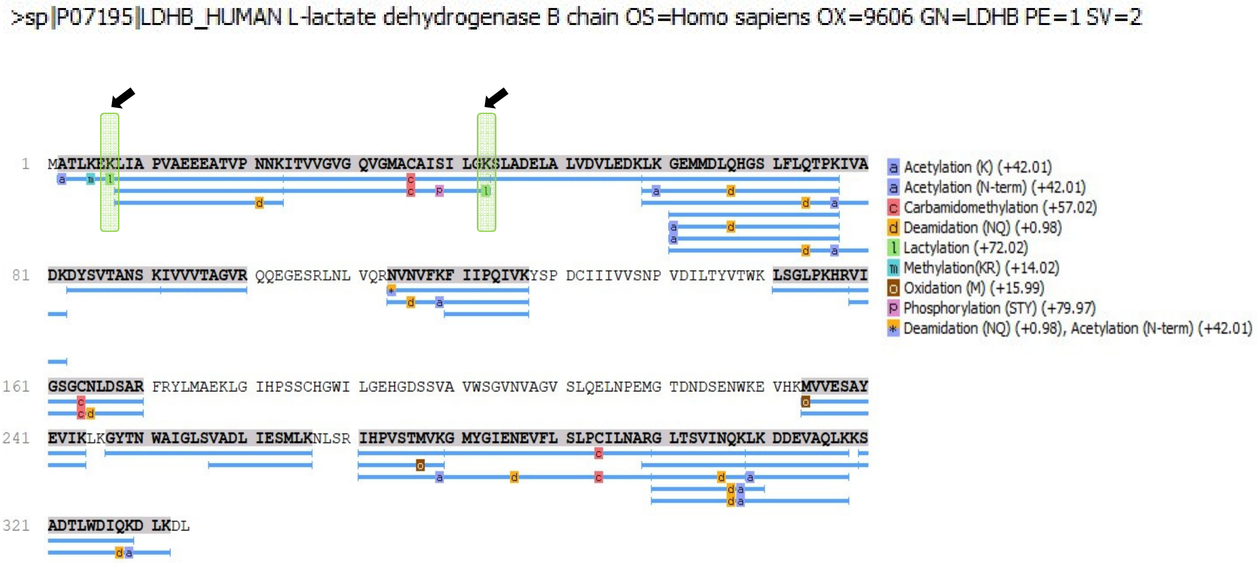

**Fig. S8v. Protein sequence coverage of human L-lactate dehydrogenase B chain (LDHB).** The amino acid sequence of human LDHB is shown. Blue lines represent peptides obtained from a HEK293 protein digest. Post-translational modifications are indicated, were lactylation is highlighted.

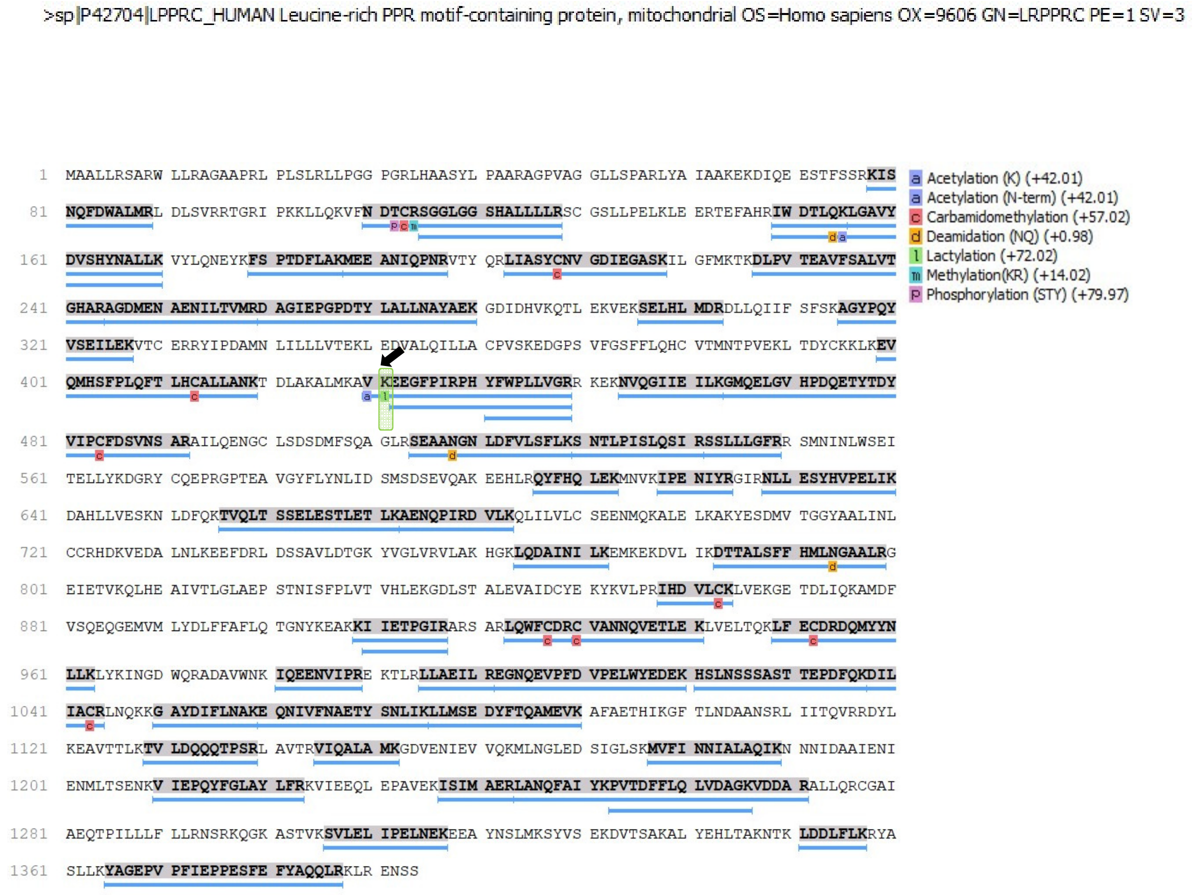


**Fig. S8vi. Protein sequence coverage of human mitochondrial Leucine-rich PPR motif-containing protein (LPPRC)**. The amino acid sequence of human LPPRC is shown. Blue lines represent peptides obtained from a HEK293 protein digest. Post-translational modifications are indicated, were lactylation is highlighted.

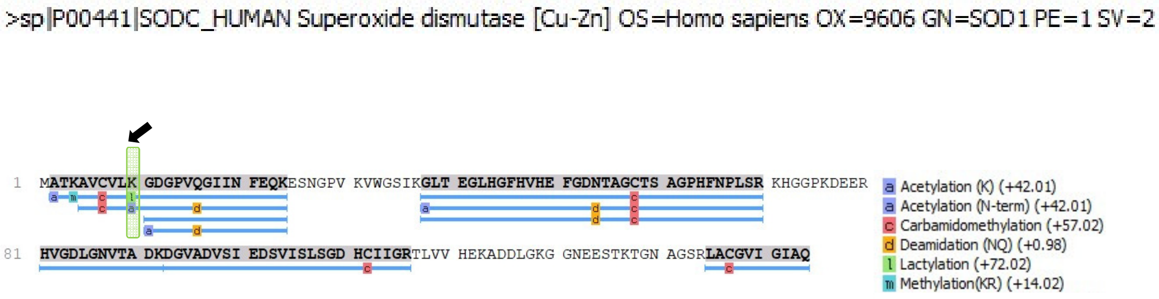


**Fig. S8vii. Protein sequence coverage of human superoxide dismutase [Cu-Zn] (SODC).** The amino acid sequence of human SODC is shown. Blue lines represent peptides obtained from a HEK293 protein digest. Post-translational modifications are indicated, were lactylation is highlighted.

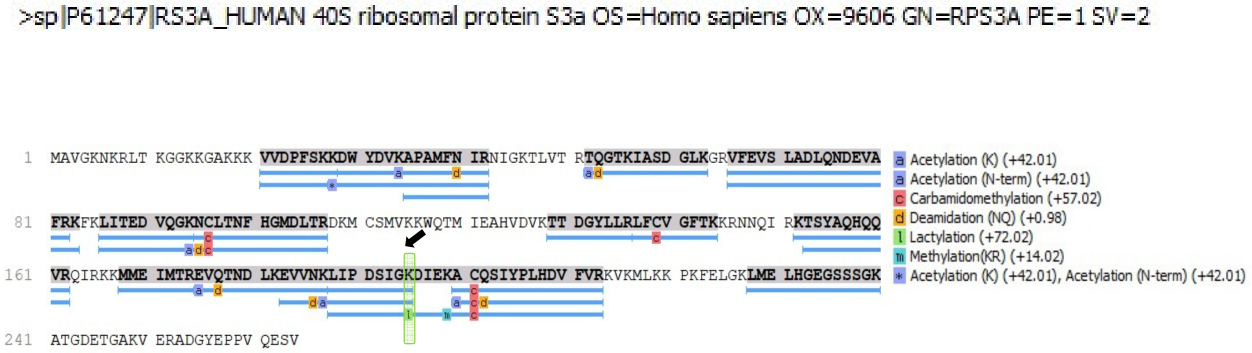


**Fig. S8viii. Protein sequence coverage of human small ribosomal subunit protein uS3 (RS3).** The amino acid sequence of human RS3 is shown. Blue lines represent peptides obtained from a HEK293 protein digest. Post-translational modifications are indicated, were lactylation is highlighted.

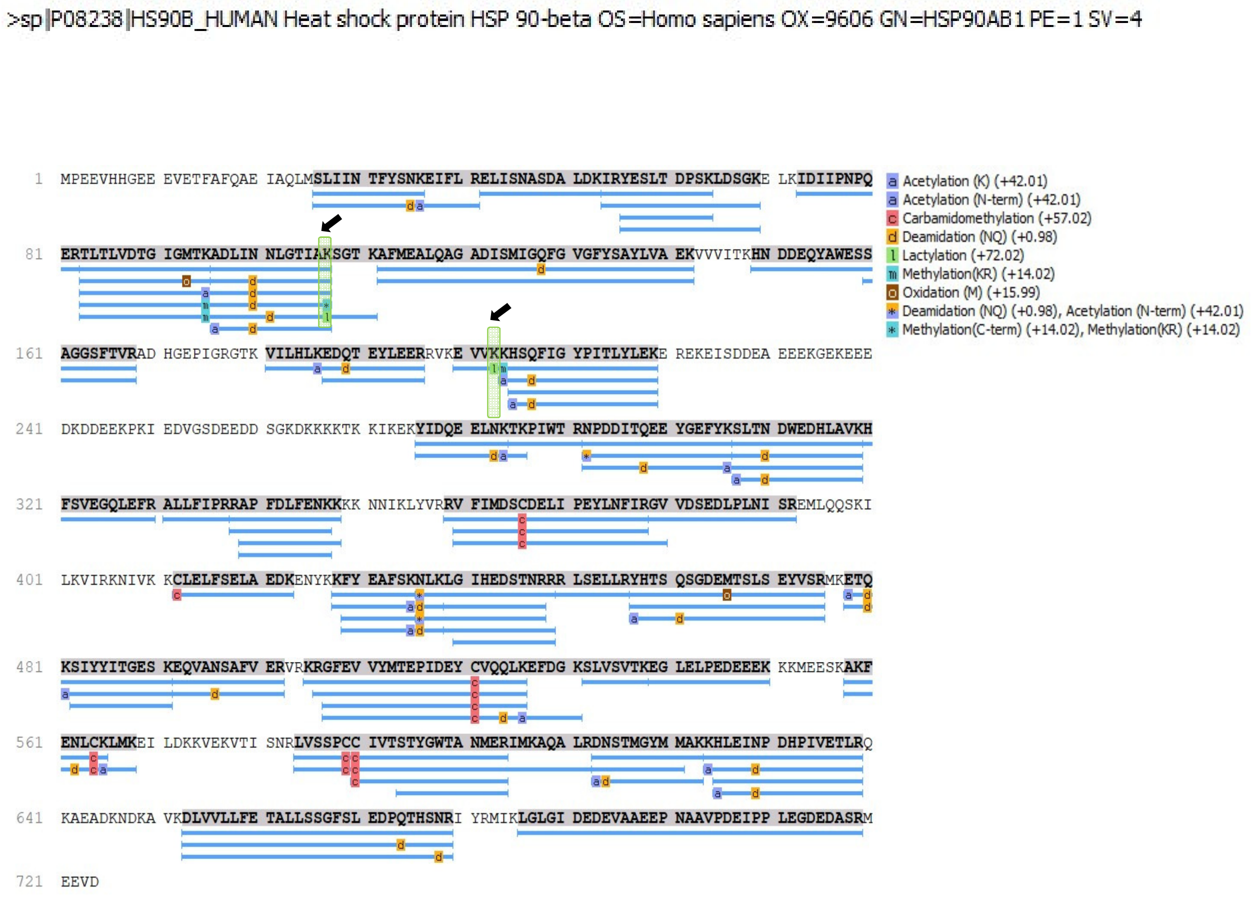


**Fig. S8ix. Protein sequence coverage of human heat shock protein HSP 90-alpha (HS90A).** The amino acid sequence of human HS90A is shown. Blue lines represent peptides obtained from a HEK293 protein digest. Post-translational modifications are indicated, were lactylation is highlighted.

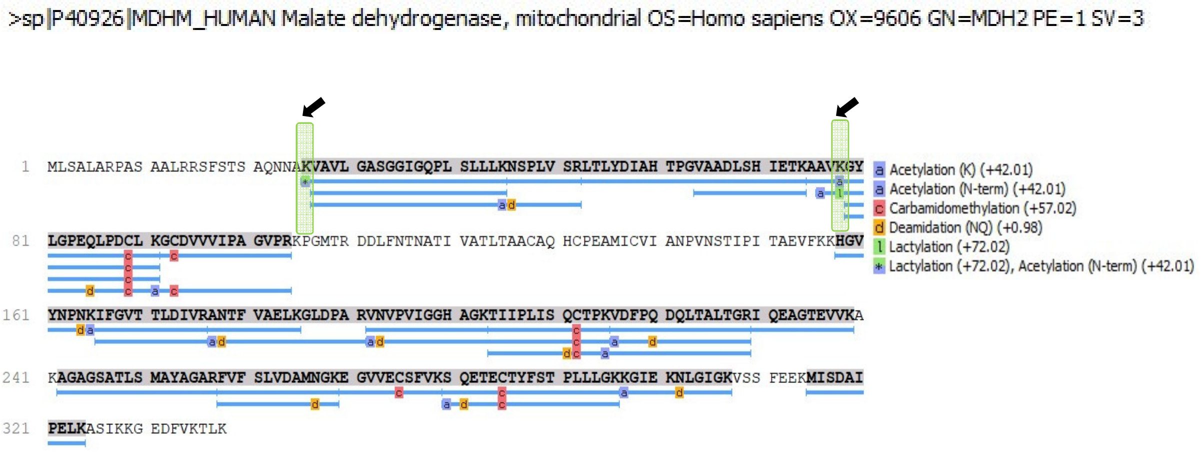


**Fig. S8x. Protein sequence coverage of human mitochondrial malate dehydrogenase (MDH2).** The amino acid sequence of human MDHM is shown. Blue lines represent peptides obtained from a HEK293 protein digest. Post-translational modifications are indicated, were lactylation is highlighted.

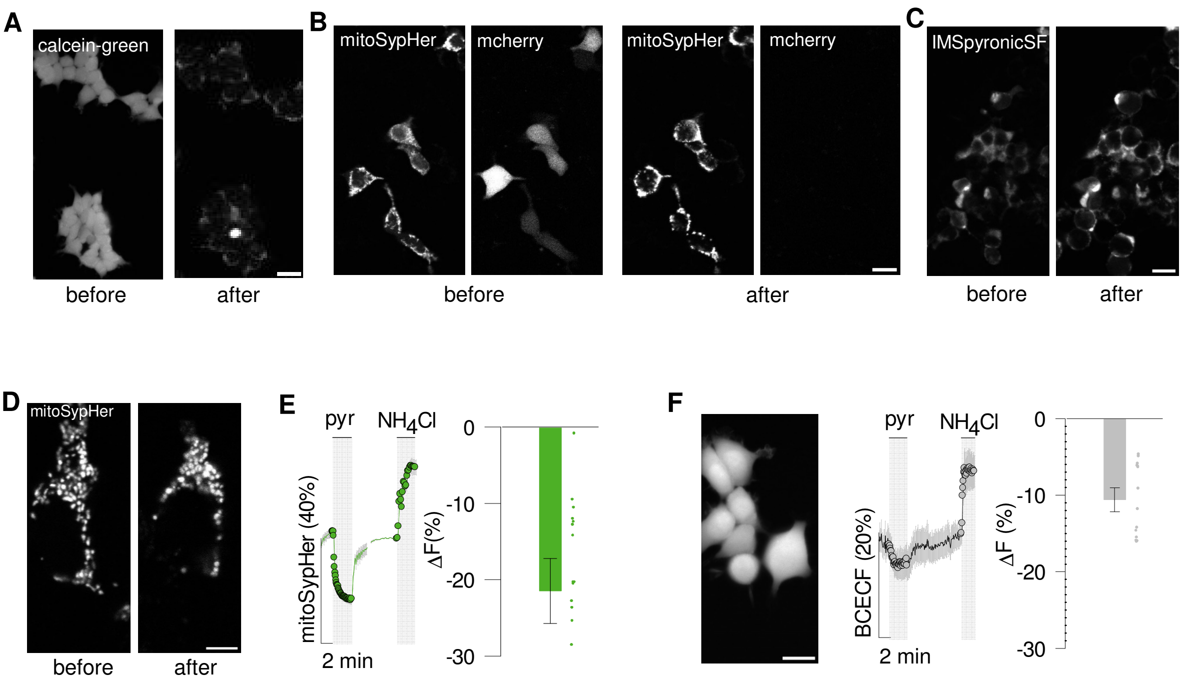


**Fig. S9. Digitonin permeabilizes the plasma membrane without apparent damage to mitochondria**. HEK293 cultures were imaged before and after a three-minute exposure to digitonin in an intracellular buffer (see Materials and Methods). All bars represent 20 µm. (**A**) Cells loaded with calcein-green, which distributes in cytosol and nucleous. Right panel, DIC image. (**B**) Cells co-expressing mitoSypHer and cytosolic mCherry. (**C**) Cells expressing pyronicSF targeted to the IMS (mitochondrial intermembrane space). (**D**) MitoSypHer distribution at higher resolution. (**E**) Effect of 10 mM pyruvate and 5 mM $NH_4Cl$ on mitochondrial pH in intact cells. Bar graph summarizes data from four experiments, 19 cells. (**F**) Effect of 10 mM pyruvate on cytosolic pH in intact cells. Bar graph summarizes data from three experiments, 14 cells.

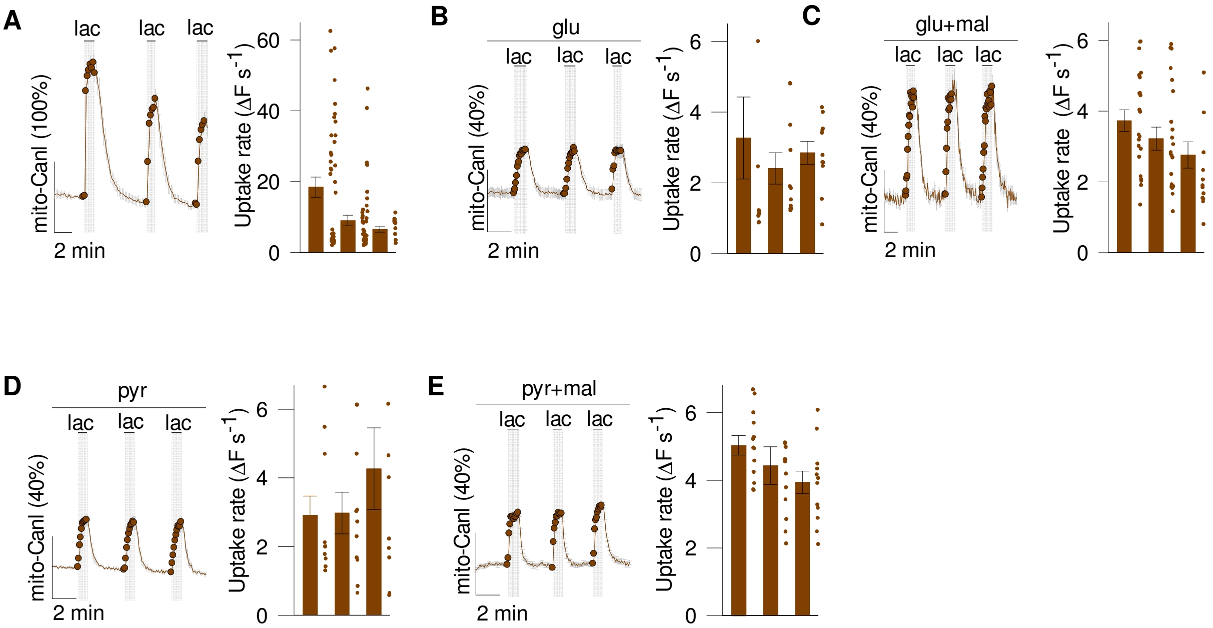


**Fig. S10. Estimation of the lactate transport rate in energized mitochondria is reproducible.** Permeabilized HEK293 cultures were exposed to repetitive lactate pulses of 1 mM (A, B) or 0.5 mM (C, D, E) in the absence or presence of mitochondrial substrates (mean ± SEM). (**A**) Without substrates (five experiments, 42 cells). (**B**) Glutamate (0.2 mM; two experiments, 11 cells). (**C**) Glutamate (0.2 mM) and malate (0.1 mM; four experiments, 22 cells). (**D**) Pyruvate (0.03 mM or 0.3 mM; two experiments, 10 cells). (**E**) Pyruvate (0.03 mM) and malate (0.1 mM; two experiments, 13 cells).

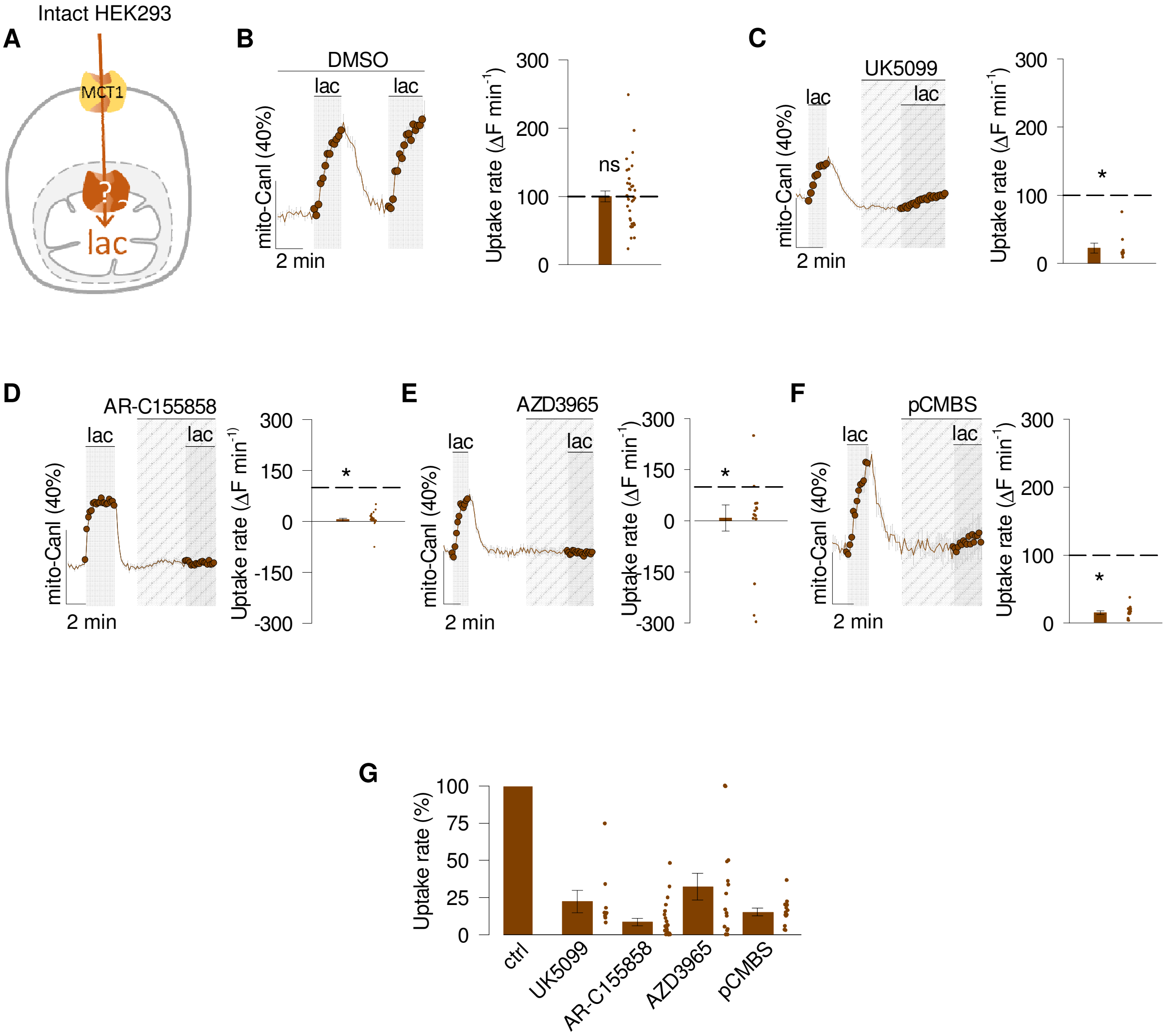


**Fig. S11. Validation of lactate transport blockers in intact cells**. Whole HEK293 cultures that expresses mito-CanlonicSF were exposed to 1 mM lactate. (**A**) Illustration of the stop-protocol. (**B**) Ctrl (DMSO 0.01-0.05%; four experiments, 35 cells). (**C**) 1 µM UK (two experiments, 9 cells). (**D**) 1µM AR (two experiments, 24 cells). (**E**) 10µM AZD (two experiments, 15 cells). **(F)** 250µM pCMBS (three experiments, 13 cells). **(G)** The bar graph summarizes the inhibitory effect.

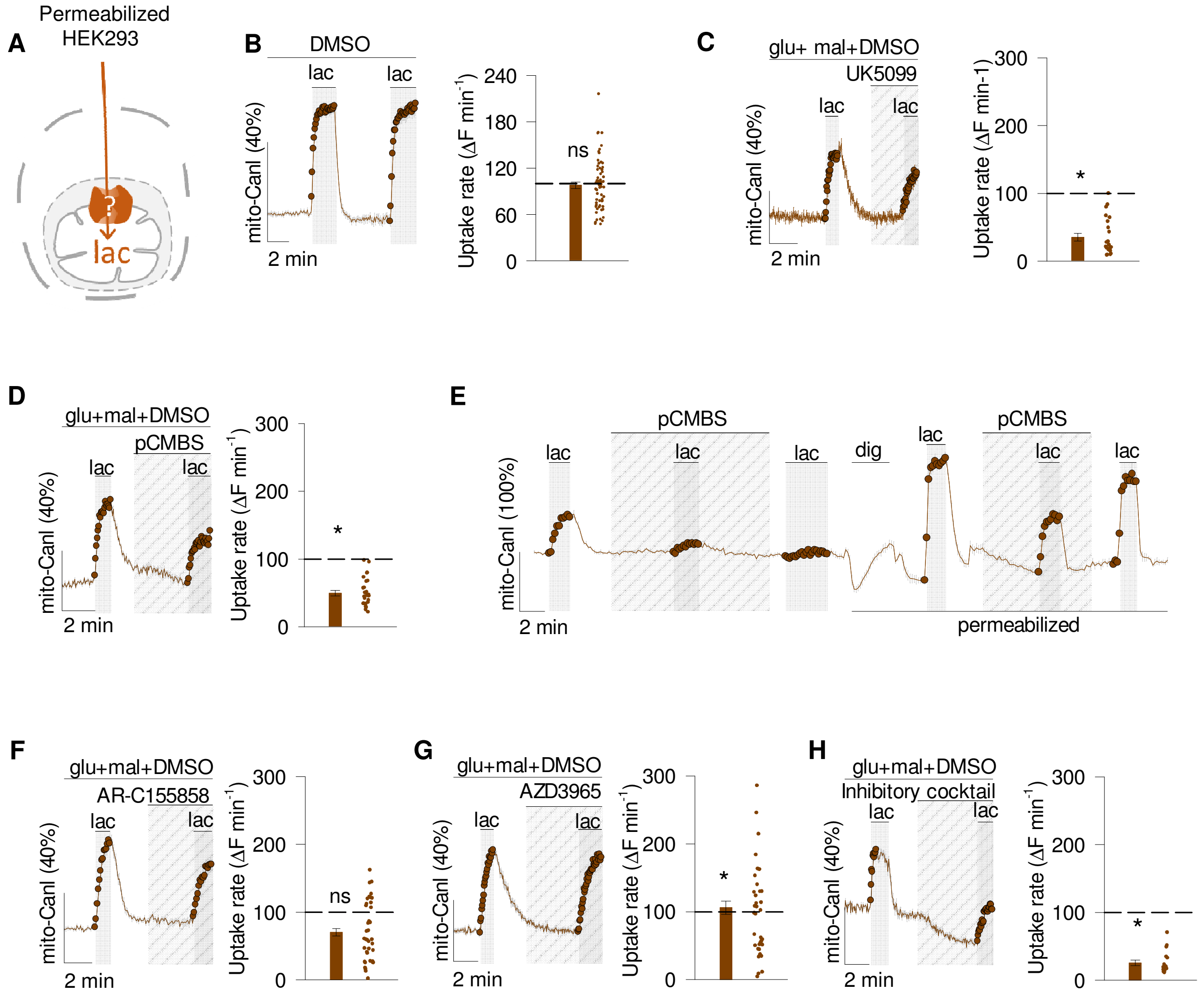


**Fig. S12. Partial inhibition of lactate transport in permeabilized cells**. Digitonin-treated HEK293 cultures expressing mito-CanlonicSF were exposed to 0.5 mM lactate under energization (0.2 mM glutamate and 0.1 mM malate). (**A**) Illustration of the inhibitory protocol. (**B**) Ctrl (DMSO 0.01%-0.05%; nine experiments, 57 cells). (**C**) UK5099 (0.5 μM; four experiments, 23 cells). (**D**) pCMBS (50 μM; four experiments, 25 cells). (**E**) Cells were exposed to 1 mM lactate and 250 μM of pCMBS before and after permeabilization with digitonin (dig)(three experiments, 11 cells). (**F**) AR-C155858 (1 μM; five experiments, 38 cells). (**G**) AZD3965 (10 μM; five experiments, 36 cells). (**H**) Inhibitor cocktail contains 1 μM UK5099, 1 μM AR-C155858, 50 μM pCMBS and 5 μM syrosingopine (three experiments, 18 cells). *, $p < 0.05$; NS, not significant, paired t-test.

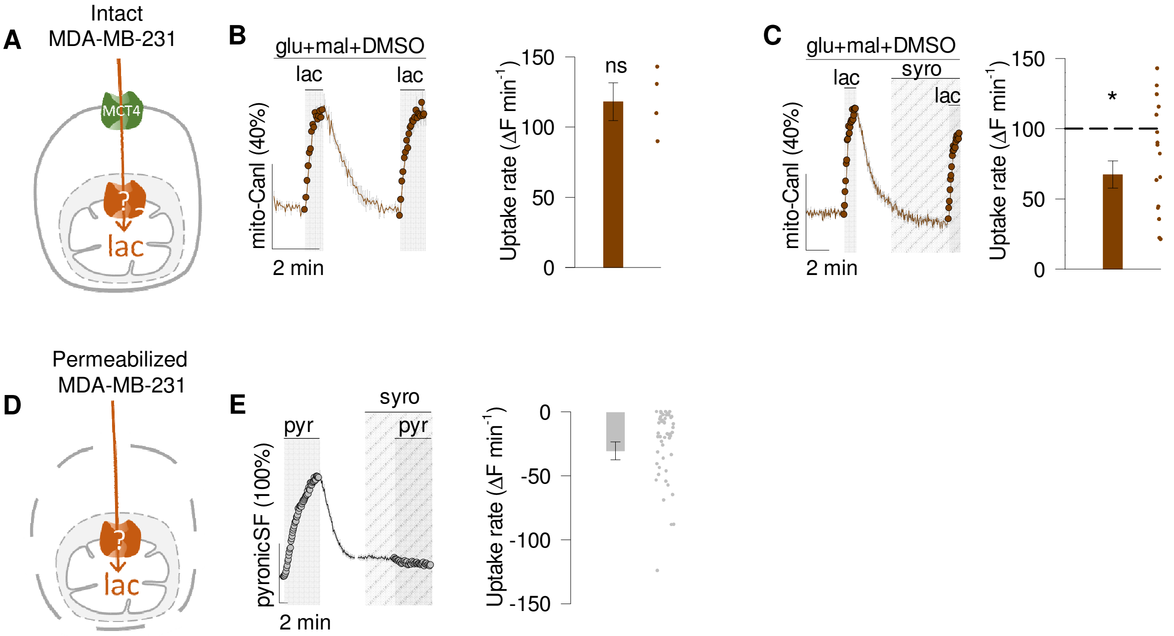


**Fig. S13. Differential effect of syrosingopine on permeabilized and intact MDA-MB-231 cells**. Cells were exposed to 0.5 lactate under incubation of 0.2mM glutamate + 0.1mM malate. (**A**) Illustration of the stop-protocol. (**B**) Ctrl (DMSO 0.01%, four cells). (**C**) 5 µM syrosingopine (two experiments, 13 cells; paired-t-test). (**D**) Illustration of the inhibitor-stop protocol. (**E**) Cells expressing PyronicSF at the cytosol with 10 mM pyruvate and 5 µM syrosingopine (four experiments, 50 cells; Wilcoxon Signed Rank Test). *; p<0.05; NS, not significant.

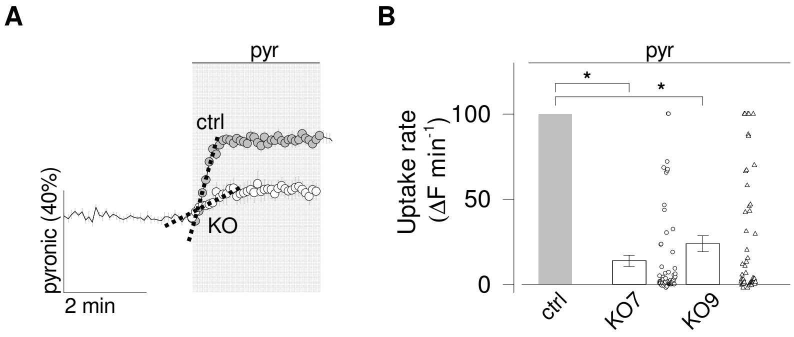


**Fig. S14. MPC2KO inhibits the initial rate of pyruvate uptake**. (**A**) Response to 10 mM pyruvate of HEK293 wild type (●) and MPC2KO (○) cells expressing pyronic in the mitochondrial matrix. Initial slopes of uptake are indicated by the dotted lines. (**B**) Summary of five wild type experiments (ctrl; 58 cells), five sgMPC2KO7 experiments (61 cells) and four sgMPC2KO9 experiments (54 cells). Wilcoxon Signed Rank Test. *; $p<0.05$.

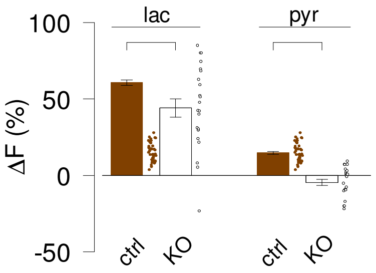


**Fig. S15. MPC2KO decreases the steady-state of lactate.** Permeabilized HEK293 cells expressing mitoCanlonicSF were targeted for MPC2 subunit knockout. Cells were energized with 0.2 mM glutamate/0.1 mM malate/ 0.1 mM pyruvate. Effect of 10 mM lactate and 10 mM pyruvate in controls (three experiments, 40 cells) and HEK293 MPC2KO cells (three experiments, 27 cells). *, $p < 0.05$; NS, not significant, paired t-test.

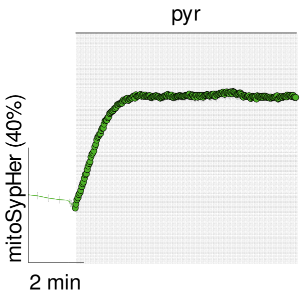


**Fig. S16. Stability of mitochondrial energization by pyruvate**. Permeabilized HEK293 cultures expressing mito-SypHer were exposed to 10 mM pyruvate. Representative of three experiments (13 cells).

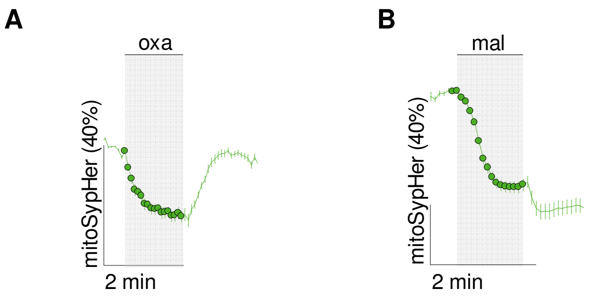


**Fig. S17. Effect of oxamate and malate on matrix pH**. HEK293 cells expressing mitoSypHer. (**A**) Cells were exposed to 10 mM lactate (two experiments, 21 cells). (**B**) Cells were exposed to 10 mM malate (three experiments, 23 cells)

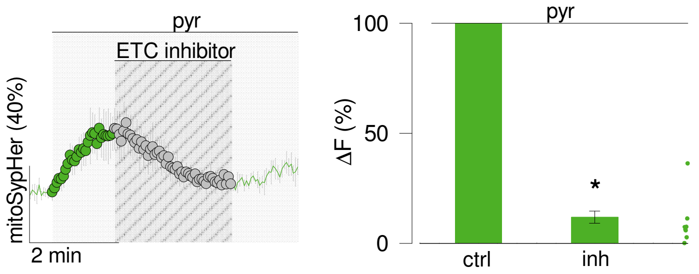


**Fig. S18. Effect of ETC inhibitors in energized neurons.** Mitochondria were exposed to pyruvate (three experiment, 13 cells) in the absence and presence of ETC blockers: rotenone (1 µM) and antimycin (1 µM), paired t-test.

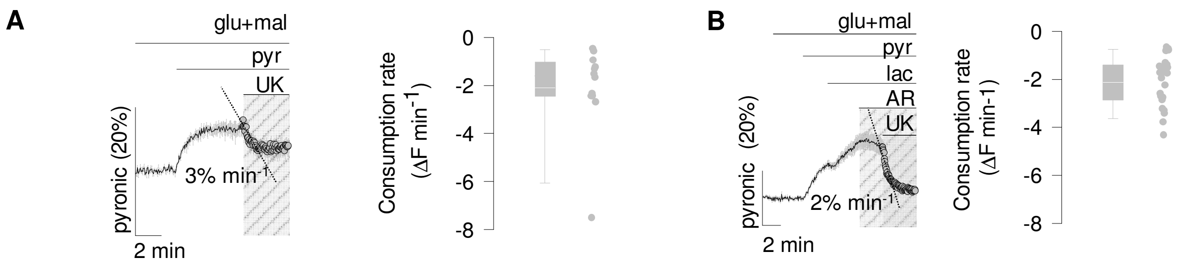


**Fig. S19. Mitochondrial pyruvate consumption.** Permeabilized HEK293 cells expressing a pyruvate sensor (pyronic) in the matrix (San Martin et la., 2014). Cells were incubated in 0.5 mM pyruvate by a transport–stop protocol under 0.2 mM glutamate and 0.1 mM malate. (**A**) 1 µM UK5099 (three experiments, 12 cells). (**B**) Cells were exposed to high lactate (10mM), 1 µM UK5099 and 1 µM AR-C155858 (four experiments, 23 cells).

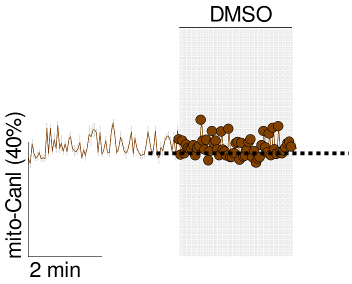


**Fig. S20. No effect DMSO on the lactate steady-state.** HEK293 cells expressing mitoCanlonicSF were exposed to 0.01% of DMSO. Data from three controls experiments (17 cells)**.**

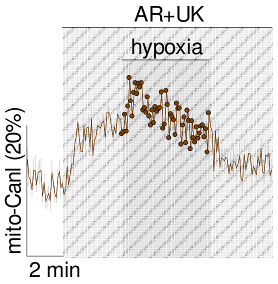


**Fig. S21. Transient accumulation of lactate by hypoxia.** Neuronal mitochondria were exposed to nitrogen-gassed buffer with 1 µM AR-C155858 and 1 µM UK5099.

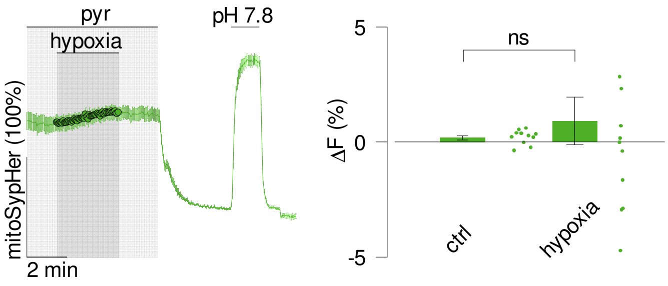


**Fig. S22. Hypoxia does not affect matrix pH.** Permeabilized neurons expressing mito-SypHer were energized with 0.1 mM pyruvate and exposed to nitrogen-gassed buffer (hypoxia). Summary of two experiments (11 cells) before and after 1 minute of hypoxia exposure.
